# Tetherin antagonism by SARS-CoV-2 enhances virus release: multiple mechanisms including ORF3a-mediated defective retrograde traffic

**DOI:** 10.1101/2021.01.06.425396

**Authors:** Hazel Stewart, Roberta Palmulli, Kristoffer H. Johansen, Naomi McGovern, Ola M. Shehata, George W. Carnell, Hannah K. Jackson, Jin S. Lee, Jonathan C. Brown, Thomas Burgoyne, Jonathan L. Heeney, Klaus Okkenhaug, Andrew E. Firth, Andrew A. Peden, James R. Edgar

**Affiliations:** Department of Pathology, University of Cambridge, Cambridge. CB2 1QP. UK; Laboratory of Immune Systems Biology, National Institute of Allergy and Infectious Diseases, National Institutes of Health, Bethesda, USA.; Department of Biomedical Science, University of Sheffield, Firth Court, Sheffield. S10 2TN. UK.; Department of Veterinary Medicine, University of Cambridge, Cambridge. CB3 0ES. UK.; Department of Infectious Disease, Imperial College London, London. W2 1PG. UK.; Royal Brompton Hospital, Guy’s and St Thomas’ NHS Foundation Trust, London. SW3 6NP. UK.; UCL Institute of Ophthalmology, University College London, London. EC1V 9EL. UK.

**Keywords:** Tetherin, SARS-CoV-2, restriction factor, ORF3a, double membrane vesicles, spike.

## Abstract

The antiviral restriction factor, tetherin, blocks the release of several different families of enveloped viruses, including the *Coronaviridae*. Tetherin is an interferon-induced protein that forms parallel homodimers between the host cell and viral particles, linking viruses to the surface of infected cells and inhibiting their release. We demonstrated that SARS-CoV-2 infection causes tetherin downregulation, and that tetherin depletion from cells enhances SARS-CoV-2 viral titres. We investigated the potential viral proteins involved in abrogating tetherin function and found that SARS- CoV-2 ORF3a reduces tetherin localisation within biosynthetic organelles via reduced retrograde recycling and increases tetherin localisation to late endocytic organelles. By removing tetherin from the Coronavirus budding compartments, ORF3a enhances virus release. We also found expression of Spike protein caused a reduction in cellular tetherin levels. Our results confirm that tetherin acts as a host restriction factor for SARS-CoV-2 and highlight the multiple distinct mechanisms by which SARS-CoV-2 subverts tetherin function.

**Author Summary:** Since it was identified in 2019, SARS-CoV-2 has displayed voracious transmissibility which has resulted in rapid spread of the virus and a global pandemic. SARS-CoV-2 is a member of the *Coronaviridae* family whose members are encapsulated by a host-derived protective membrane shell. Whilst the viral envelope may provide protection for the virus, it also provides an opportunity for the host cell to restrict the virus and stop it spreading. The anti-viral restriction factor, tetherin, acts to crosslink viruses to the surface of infected cells and prevent their spread to uninfected cells. Here, we demonstrate that SARS-CoV-2 undergoes viral restriction by tetherin, and that SARS-CoV-2 moves tetherin away from the site of Coronavirus budding to enhance its ability to escape and infect naïve cells. Tetherin depletion from cells enhanced SARS-CoV-2 viral release and increased propagation of the virus. We found that the SARS-CoV-2 protein, ORF3a, redirects tetherin away from the biosynthetic organelles where tetherin would become incorporated to newly forming SARS-CoV-2 virions – and instead relocalises tetherin to late endocytic organelles. We also found that SARS-CoV-2 Spike downregulates tetherin. These two mechanisms, in addition to the well described antagonism of interferon and subsequent ISGs highlight the multiple mechanisms by which SARS-CoV-2 abrogates tetherin function. Our study provides new insights into how SARS-CoV-2 subverts human antiviral responses and escapes from infected cells.

## Introduction

The causative agent of coronavirus disease 2019 (COVID-19) is the enveloped coronavirus, severe acute respiratory syndrome coronavirus 2 (SARS-CoV-2) [1, 2]. Sarbecoviruses are positive-sense single stranded RNA viruses, and their genomes encode a large non-structural replicase complex and four main structural proteins, spike (S), membrane (M), envelope (E), and nucleocapsid (N). In addition, the SARS-CoV-2 genome encodes other accessory proteins that facilitate replication, cell entry and immune evasion.

Viruses can be broadly categorized by the presence or absence of a host-derived lipid envelope. Membrane envelopment protects the viral capsid from the external environment, reduces host immune recognition, and aids viral entry to new cells. However, this envelopment process does provide an opportunity for host cells to integrate anti-viral factors within the forming virions.

SARS-CoV-2 cellular entry is mediated by spike protein binding to the host receptor angiotensin-converting enzyme-2 (ACE2) [3]. Unlike that of 2002-2003 SARS-CoV (hereafter named SARS-CoV-1), the SARS-CoV-2 spike protein contains a polybasic furin cleavage site which facilitates the cleavage of the spike into two proteins, S1 and S2 that remain non-covalently associated [4, 5]. The S2 fragment is further primed by the serine protease TMPRSS2 [3], whilst the S1 fragment binds Neuropilin-1 [6, 7], facilitating virus entry and infection. Coronaviruses enter TMPRSS2-positive cells by direct fusion at the plasma membrane, and are endocytosed by TMPRSS2-negative cells [8], following which their envelope fuses within late endosomes/lysosomes [9], liberating the viral nucleocapsid to the cytosol of the cell. Subsequently, viral proteins are translated and assembled at modified tubulovesicular ERGIC (endoplasmic reticulum-Golgi intermediate compartment) organelles. Coronaviruses modify host organelles to generate viral replication factories, so-called DMVs (double-membrane vesicles) that act as hubs for viral RNA synthesis [10].

Tetherin (*Bst2*) is an interferon-inducible restriction factor that limits egress of several types of enveloped viruses, reducing the infection of neighbouring naïve cells. Tetherin is a type II integral membrane protein with a short cytosolic tail, a single pass transmembrane domain, and an extracellular coiled-coil domain that is anchored to the membrane via a C terminal GPI anchor. Cysteine residues in the extracellular domain mediate homodimer formation via disulphide bridges, linking tetherin molecules on virus and cell membrane, leading to the retention of nascent viral particles on the surface of infected cells [11]. The cell surface retention of virions enhances their reinternalization to endosomal compartments, limiting the extent of virus spread [12]. Tetherin is a type-I Interferon-Stimulated Gene (ISG), and so although it is constitutively expressed by many cell types, expression of tetherin is enhanced by the presence of type-I interferon (IFN) [13].

Enveloped viruses including human immunodeficiency virus 1 (genus *Lentivirus*) [13, 14], Ebola virus (genus *Ebolavirus*) [13, 15], Kaposi’s sarcoma-associated herpesvirus (KSHV) (genus *Rhadinovirus*) [16] and human coronavirus 229E (HCoV- 229E) (*Alphacoronavirus*) [17] undergo tetherin-dependent restriction. For enveloped viruses to produce fully released progeny, they have evolved means to counteract tetherin activity. Although the molecular mechanisms by which each virus downregulates tetherin differ, they all reduce tetherin abundance from the organelle in which respective viruses bud, and/or reduce tetherin levels upon the plasma membrane.

Of the previously described coronaviruses, HCoV-229E and SARS-CoV-1 have been shown to undergo viral restriction by tetherin [17, 18]. Two SARS-CoV-1 proteins have been shown to antagonise tetherin resulting in a concomitant increase in virion spread – the ORF7a protein and spike glycoprotein [18, 19]. However, several questions remain about the mechanisms surrounding tetherin antagonism by coronaviruses. It is also unclear exactly how and where tetherin forms such tethers, as in coronaviruses, unlike other viruses that undergo tetherin-dependent restriction (such as HIV-1), membrane envelopment occurs in the biosynthetic pathway and not at the plasma membrane. The mechanism by which sarbecoviruses dysregulate tetherin remain unclear.

Here, we show that tetherin is directly responsible for tethering of nascent enveloped SARS-CoV-2 virions to infected cell surfaces. We demonstrate using primary cells and immortalised cell lines that SARS-CoV-2 infection causes a dramatic downregulation of tetherin, and that loss of tetherin aids SARS-CoV-2 viral spread and infection. We examine the effect of expression of individual SARS-CoV-2 proteins on tetherin antagonism and find that ORF3a redirects tetherin away from the biosynthetic pathway and to the endolysosomal pathway by inhibiting retrograde retrieval. The reduction of tetherin within the biosynthetic pathway limits its incorportation to forming virions, with subsequent enhancement in virus release. We also demonstrate that Spike expression causes tetherin downregulation, as has previously been described for SARS-CoV-1.

## Results

### SARS-CoV-2 downregulates tetherin

Airway epithelial cells are one of the initial sites of viral contact during SARS-CoV-2 infection. Differentiation of nasal primary human airway epithelial (HAE) cells results in a pseudostratified epithelium consisting of ciliated, goblet and basal cells, and SARS-CoV-2 readily infects ciliated cells [20]. Differentiated, mock-infected HAE cells displayed negligible levels of tetherin (**Supplemental Figure 1A**), reflecting low basal tetherin expression in the absence of IFN stimulation. Differentiated HAE cells were infected with SARS-CoV-2 and sections analysed by immunofluorescence microscopy to detect changes in tetherin. Infected cells, confirmed by anti-spike labelling, displayed lower tetherin fluorescence than neighbouring, uninfected cells (**Figure 1A, Supplemental Figure 1B**). The levels of tetherin in mock-infected HAE cells is lower than observed in uninfected neighbours in infected wells. This bystander effect indicates that infected cells are still able to generate IFN to act upon uninfected neighbouring cells, enhancing tetherin expression.

**Figure 1.**
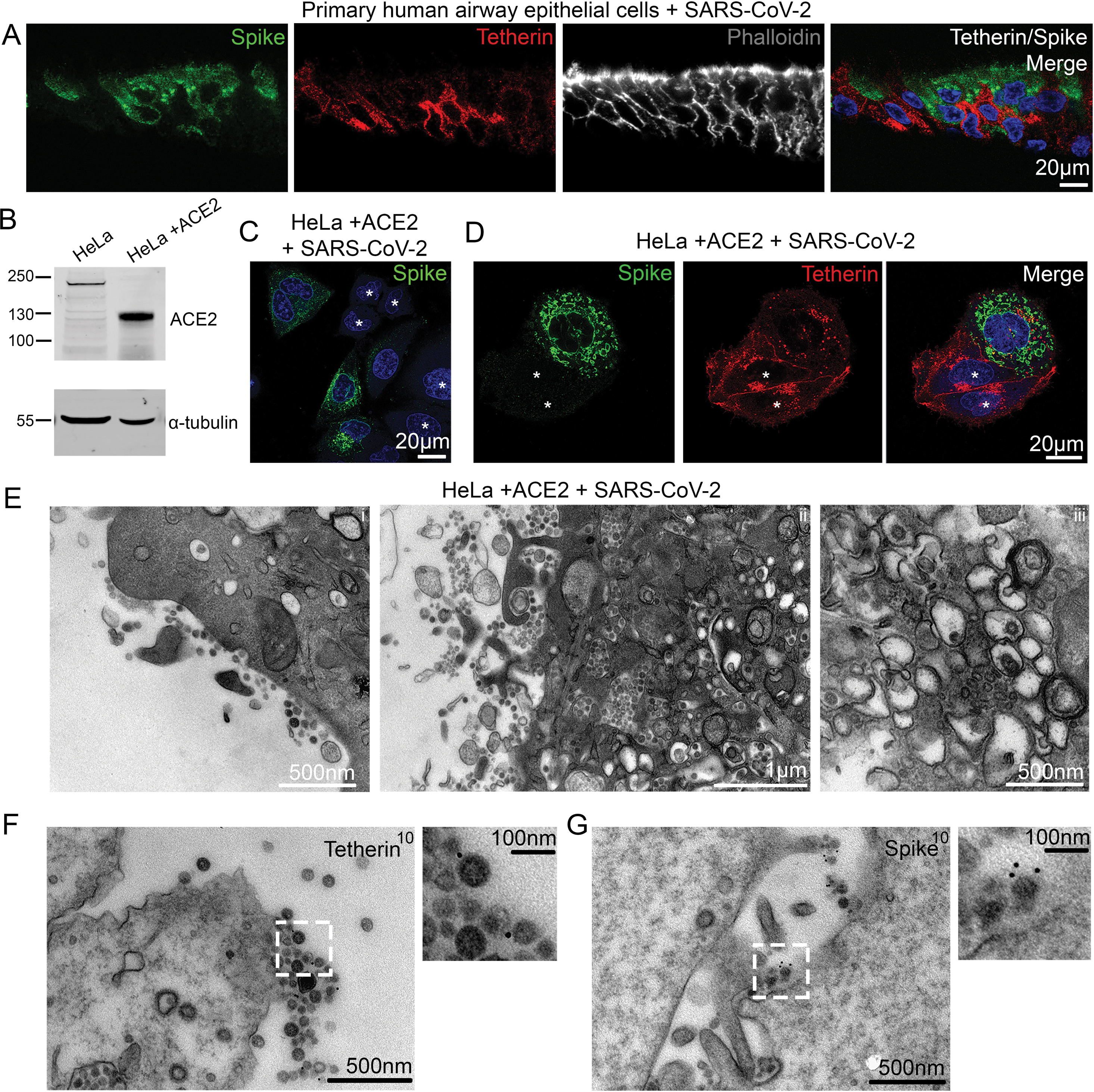
SARS-CoV-2 infection downregulates tetherin in primary human airway epithelial and HeLa+ACE2 cells. (A) Nasal primary human airway epithelial (HAE) cells were infected with SARS- CoV-2 (MOI 0.01). Cells were fixed at 48 hpi and embedded to OCT. Cryostat sections were collected and prepared for confocal microscopy. Sections were immunolabelled with antibodies against SARS-CoV-2 spike (green) - to reveal SARS-CoV-2 infected cells, tetherin (red), and with phalloidin-647 (grey) and DAPI (blue). (B) ACE2 expressing stable HeLa cell lines were generated by lentiviral transduction. To confirm ACE2 expression, mock and ACE2 transduced cells were lysed and immunoblotted with anti-ACE2 antibodies. Anti-tubulin was used as a loading control. (C) HeLa+ACE2 cells were infected with SARS-CoV-2 (MOI 0.5). Cells were fixed at 24 hpi and stained with antibodies against SARS-CoV-2 spike (green), and with DAPI (blue). (D) HeLa+ACE2 cells were infected with SARS-CoV-2 (MOI 0.5). Cells were fixed at 24 hpi and stained with antibodies against SARS-CoV-2 spike (green), tetherin (red), and with DAPI (blue). Uninfected cells are shown by asterisks. (E) Electron micrographs showing plasma membrane-associated SARS-CoV-2 virions, and virus filled intracellular organelles. SARS-CoV-2 infected HeLa+ACE2 cells (MOI 0.5) were fixed at 24 hpi and processed for TEM. Micrographs show (i) plasma membrane-associated virions, (ii) virus-filled tubulovesicular compartments directed towards the plasma membrane, (iii) virions within DMVs. (F) Surface immunogold electron microscopy of SARS-CoV-2 infected HeLa+ACE2 cells. Cells were infected with SARS-CoV-2 (MOI 0.5), fixed at 24 hpi and immunogold surface labelled using an anti-tetherin antibody. (G) as in (F) but cells were labelled with an anti-SARS-CoV-2 spike antibody.

A robust IFN response is considered a key first line of defence against viral infection. Coronaviruses have developed multiple strategies to dampen IFN responses, and a hallmark of SARS-CoV-2 infection is the very weak IFN response [21]. As an IFN stimulated gene, tetherin transcription is likely antagonised by SARS-CoV-2. To determine whether SARS-CoV-2 reduces tetherin independently of IFN antagonism, we assessed the impact of SARS-CoV-2 infection on HeLa cells which constitutively express high levels of tetherin, without the need of induction through IFN. To allow SARS-CoV-2 infection, we generated a HeLa cell line stably expressing ACE2 by lentiviral transduction, designated as HeLa+ACE2, and ACE2 protein expression was confirmed by Western blotting (**Figure 1B**). Using a clinical isolate of SARS- CoV-2 (isolate BetaCoV/Australia/VIC01/2020) [22], we performed viral infection assays and fixed the cells 24 hours post infection (hpi). Infected cells were confirmed by anti-spike labelling (**Figure 1C**, uninfected cells shown with asterisk).

Uninfected HeLa+ACE2 cells display tetherin localised to the plasma membrane, perinuclear and some more peripheral punctate organelles. Upon SARS-CoV-2 infection, tetherin signal was lost from the plasma membrane and perinuclear region, (**Figure 1D, Supplemental Figure 1C**, uninfected cells shown with asterisk). While tetherin appeared to be broadly downregulated from the plasma membrane, some plasma membrane staining remained in these cells that often colocalised with spike labelling (**Supplemental Figure 1C**), likely representing areas of surface-tethered SARS-CoV-2 virus.

To examine whether SARS-CoV-2 virions were tethered to the cell surface, we performed transmission electron microscopy. In infected HeLa+ACE2 cells, SARS- CoV-2 virions could be found clustered on the plasma membrane of cells, although tethered virions were frequently polarised to discrete areas, rather than distributed evenly along the plasma membrane (**Figure 1E, Supplemental Figure 1C**). Virus- containing tubulovesicular organelles were often polarised towards sites of significant surface-associated virus.

Electron microscopy also verified the presence of double membrane vesicles (DMVs) in infected cells, and typical Golgi cisternae were not present in infected cells (**Figure 1E**). To check whether the formation of DMVs and aberration to the biosynthetic machinery was causing a global downregulation of surface proteins, we stained infected cells for the surface protein beta2microglobulin (**Supplemental Figure 1D**), but no obvious loss in plasma membrane staining was observed in infected cells. Surface labelling immunogold electron microscopy (see Methods) revealed tetherin molecules to be found between SARS-CoV-2 virions (**Figure 1F**) and virions were verified as being SARS-CoV-2 using an anti-SARS-CoV-2 spike antibody (**Figure 1G**).

The human alveolar epithelial cell line, A549, expresses low levels of ACE2 endogenously, although ectopic overexpression of ACE2 has been used to facilitate betacoronavirus entry [23, 24]. A549 cells stably expressing ACE2, designated A549+ACE2, were generated by lentiviral transduction (**Figure 2A**) and these cells were amenable to SARS-CoV-2 infection (**Figure 2B**, uninfected cells shown with asterisk). A549 cells do not express tetherin at steady state, although its expression can be induced through stimulation with IFN alpha (IFNα) [13, 25] (**Supplemental Figure 2A**), following which tetherin is localised to the plasma membrane and to intracellular compartments. Immunofluorescence analysis of A549+ACE2 cells infected with SARS-CoV-2 revealed a dramatic loss of tetherin as revealed by the loss of tetherin in SARS-CoV-2 infected cells (**Figure 2C**, uninfected cells shown with asterisk). To examine what effect this near-total loss of tetherin had on virus tethering, we again performed electron microscopy. SARS-CoV-2 infected, IFNa treated A549+ACE2 cells were characterised by significant intracellular remodelling, but very few surface-associated virions were present, likely due to the significant tetherin downregulation (**Figure 2D**). Virion-containing DMVs were frequently observed in the perinuclear region of infected cells, and these were associated with dramatic membrane remodelling, including a loss of typical Golgi cisternae from cells (**Figures 2E, 2F**).

**Figure 2.**
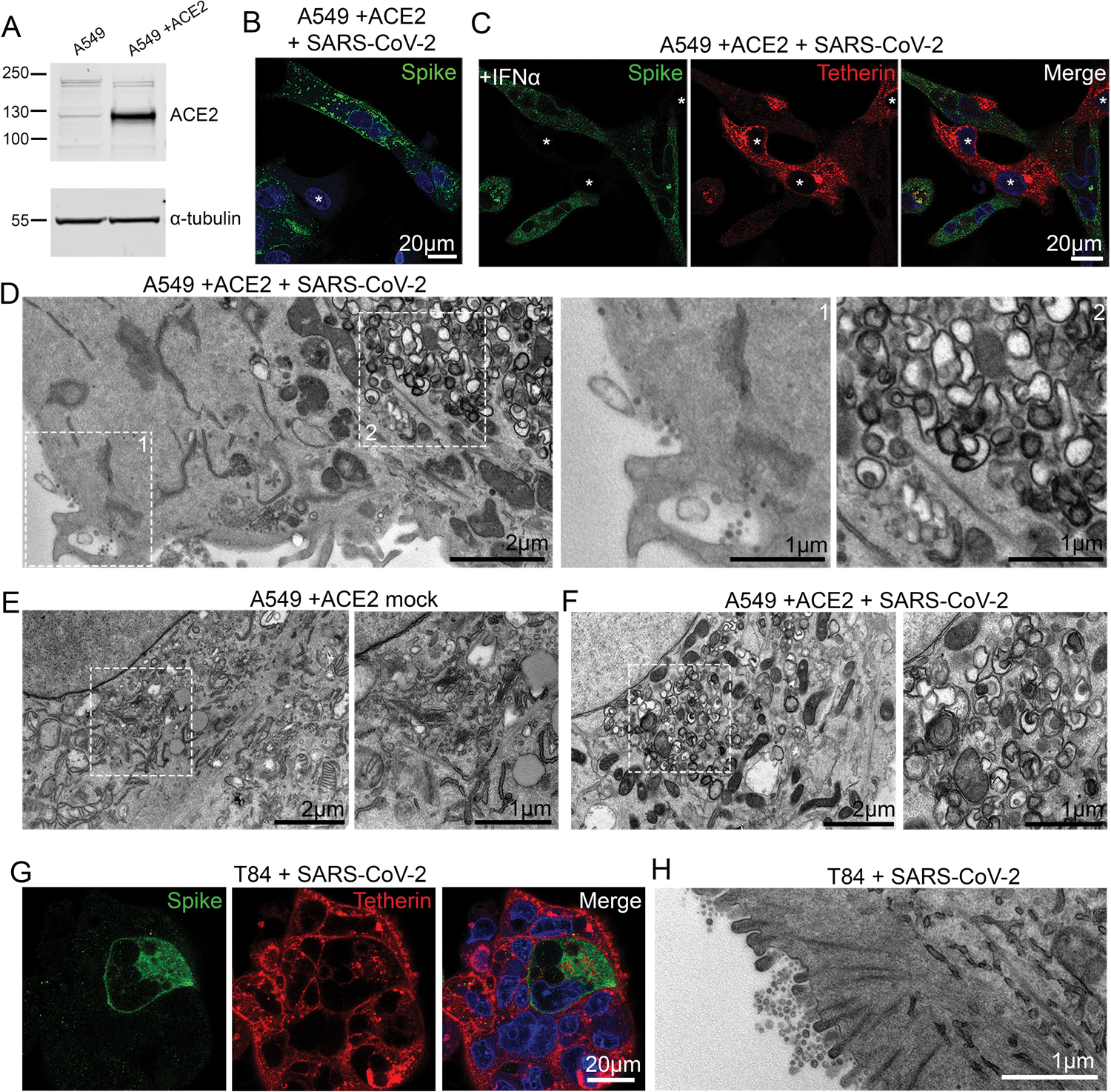
SARS-CoV-2 infection of epithelial cell lines. (A) A549 cells were transduced with ACE2 lentivirus to generate stable cell lines. Mock and ACE2 transduced cells were lysed and immunoblotted with anti- ACE2 antibodies. Anti-tubulin served as a loading control. (B) A549+ACE2 cells were infected with SARS-CoV-2 (MOI 0.5). Cells were fixed at 24 hpi and stained for SARS-CoV-2 spike (green) to reveal infected cells, and DAPI (blue). Uninfected cells are shown by asterisks. (C) A549+ACE2 cells were treated with IFNα (1000 U/mL, 24 hours) to upregulate tetherin expression. Cells were then infected with SARS-CoV-2 (MOI 0.5) and fixed at 24 hpi. Cells were immunolabelled using antibodies against SARS-CoV-2 spike (green), tetherin (red), and with DAPI (blue). Uninfected cells shown with asterisks. (D) A549+ACE2 cells were treated with IFNα (1000 U/mL) and infected with SARS-CoV-2 (MOI 0.5), fixed at 24 hpi and processed for TEM. Infected cells displayed a low frequency of virions at the plasma membrane (left inset) but significant DMV formation was observed (right inset). (E) Electron microscopy of the perinuclear region of mock A549+ACE2 cell. Zoomed area highlights typical perinuclear Golgi morphology. (F) Electron microscopy of the perinuclear region of SARS-CoV-2 infected A549+ACE2 cells. Infected cells were infected at an MOI of 0.5 and fixed at 24 hpi. Zoomed area highlights the loss of typical Golgi cisternae, and the appearance of DMVs. (G) T84 cells were infected with SARS-CoV-2 (MOI 0.5) and fixed at 24 hpi and stained for spike (green) and tetherin (red). (H) T84 cells were infected with SARS-CoV-2 (MOI 0.5) and fixed at 24 hpi and processed for TEM. Tethered virions were frequently present at the plasma membrane.

Although SARS-CoV-2 infections predominately cause pathology to the respiratory system, gastrointestinal epithelial cells, where ACE2 is highly expressed, are also infected. The human colonic adenocarcinoma epithelial cell line, T84, which express endogenous ACE2 [26] were examined for their ability to be infected by SARS-CoV- 2 and to tether virions (**Figure 2G**). Electron microscopy analysis of infected T84 cells revealed significant virus tethering at microvilli (**Figure 2H**) along flat regions of the plasma membrane, and formation of virus-filled intracellular compartments (**Supplemental Figure 2B**). The differences in the amount of tetherin downregulation between these cell lines may reflect either differences in kinetics of infection, differences in resting levels of tetherin, or differences in host machinery involved in the process of downregulation. These data are consistent with previous observations suggesting that other coronaviruses downregulate tetherin [17–19].

### Tetherin loss aids SARS-CoV-2 viral spread

To determine whether tetherin plays a functional role in viral tethering, we performed both one-step (MOI of 5) and multi-step (MOI of 1) growth curves, to measure both released and intracellular virus production from infected WT HeLa+ACE2 and Bst2KO HeLa+ACE2 (**Figure 3A**). HeLa cells were used due to their high, homogenous and IFN-independent expression of tetherin. Bst2KO HeLa cells used were previously described [27]. Both WT HeLa+ACE2 and Bst2KO HeLa+ACE2 were able to be infected with SARS-CoV-2 (**Figure 3B**). WT HeLa+ACE2 and Bst2KO HeLa+ACE2 cells were infected at the respective MOI and released and intracellular virus was harvested at the indicated time points (**Figure 3C, 3D**).

**Figure 3.**
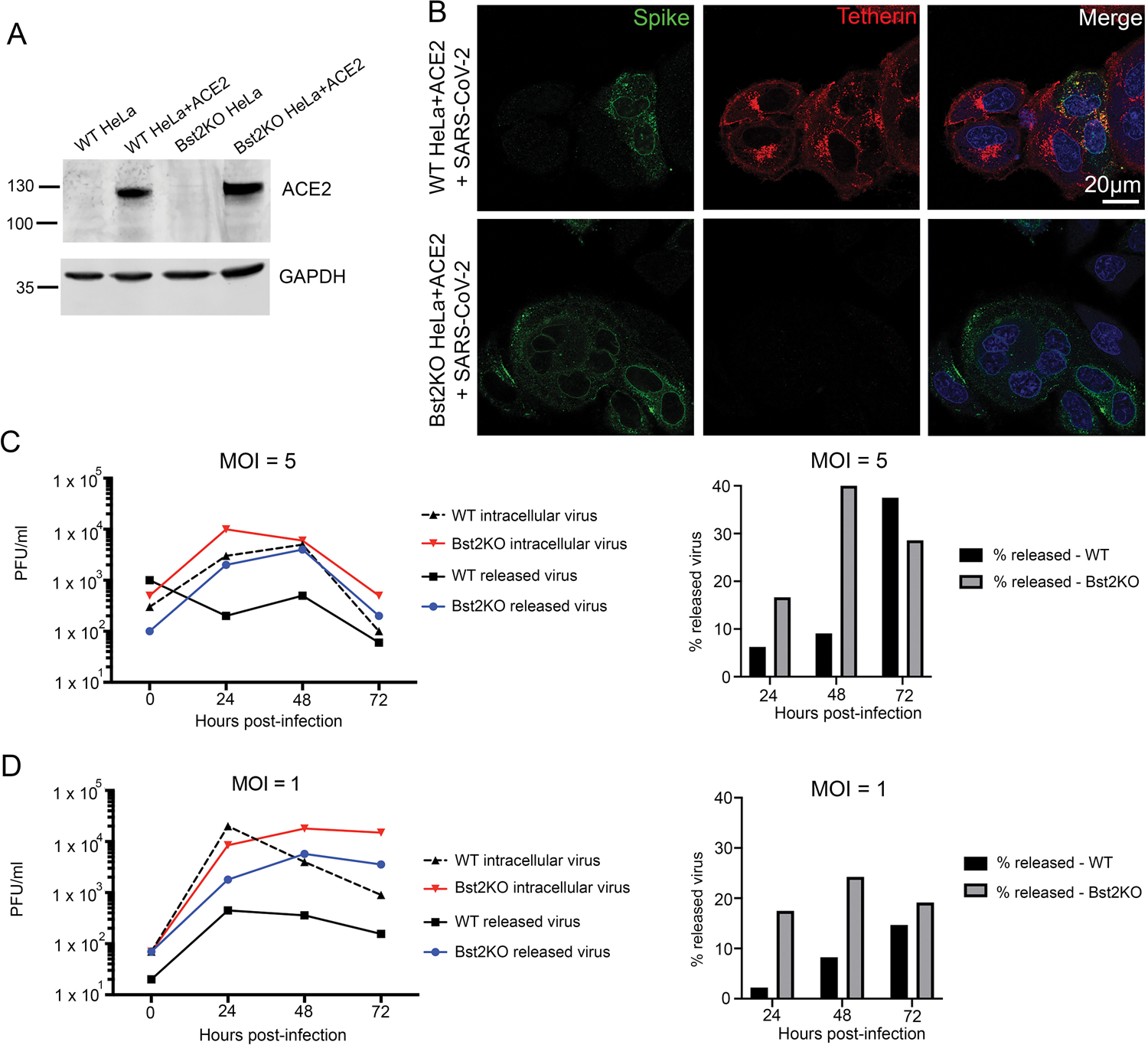
Viral growth curves reveal tetherin loss enhances viral spread. (A) Lentiviral ACE2 was used to generate stable HeLa^Bst2KO^ +ACE2 cells, and ACE2 expression was verified by Western blotting. GAPDH served as a loading control. (B) WT HeLa+ACE2 and Bst2KO HeLa+ACE2 cells were infected with SARS- CoV-2 (MOI 0.5) and fixed at 24 hpi. Cells were stained for spike (green) to demonstrate infection with SARS-CoV-2, and tetherin (red). (C) High MOI viral growth curves were performed by infecting WT HeLa+ACE2 and Bst2KO HeLa+ACE2 cells with SARS-CoV-2 at MOI (5). Released and intracellular viral titres were measured by plaque assays. (D) Low MOI viral growth curves were performed by infecting WT HeLa+ACE2 and Bst2KO HeLa+ACE2 cells with SARS-CoV-2 at MOI (1). Released and intracellular viral titres were measured by plaque assays.

The higher MOI of 5, used in the one-step growth curve (**Figure 3C**), ensures a synchronous infection event of 99.3 % of the cells (according to the Poisson distribution). This is predicted to result in synchronised RNA replication, virion assembly and egress, allowing a clear distinction between the various virus life cycle steps, as minimal reinfection should occur. However, this approach results in the vast majority of cells being infected with >1 virus particle. In addition to being less physiologically relevant, this higher viral load may be able to overcome host restriction factors and mask a phenotype. In this case, the released virus titre was clearly higher in the Bst2KO HeLa+ACE2 cells (24, 48 and 72 hpi), evidencing the tetherin-mediated restriction in the WT HeLa+ACE2 cells. However, intracellular virions appeared to accumulate quickly in the Bst2KO HeLa+ACE2 cells at the early time point of 24 hours; whether this is due to a lack of tetherin indirectly allowing enhanced RNA replication or is a by-product of this model system remains to be seen. It is obvious, however, that when the released virions are viewed as a proportion of total infectious particles, the Bst2KO HeLa+ACE2 cells release significantly more than WT HeLa+ACE2 cells at 24 and 48 hpi, due to their inability to tether nascent virions.

By contrast, the multi-step growth curve utilising an MOI of 1 (**Figure 3D**) has the advantage of being more physiologically relevant and providing a more realistic stoichiometric ratio of viral to host proteins, making it less likely to mask naturally occurring interactions and their resulting phenotypes. However as approximately 37% of the cells will not be infected by the initial inoculum, infection of naïve cells will continue to occur throughout the time course and the viral replication events cannot be presumed to be aligned. In this case, the disparity between Bst2KO HeLa+ACE2 and WT HeLa+ACE2 cells is clear until the final time point of 72 hours, with the Bst2KO HeLa+ACE2 cells continually releasing a higher proportion of virions.

Together, these data demonstrate that tetherin acts to limit SARS-CoV-2 infection and that SARS-CoV-2 acts to downregulate tetherin. These data support the notion that tetherin exerts a broad restriction against numerous enveloped viruses, regardless of whether budding occurs at the plasma membrane or within intracellular compartments. As previously studied coronaviruses, including HCoV-229E and SARS-CoV-1, have been demonstrated to downregulate tetherin [17–19] we next aimed to determine which SARS-CoV-2 protein is responsible for tetherin downregulation.

### ORF7a protein does not alter endogenous tetherin abundance, glycosylation or dimer formation

ORF7a has been shown to antagonise tetherin in both SARS-CoV-1 [18] and SARS- CoV-2 [28], although the mechanism of antagonism is not well understood. SARS- CoV-2 ORF7a has acquired a number of mutations which exist in and around the transmembrane domain which plays a role in SARS-CoV-1 ORF7a localisation. We found that SARS-CoV-1 ORF7a colocalises with the trans-Golgi marker TGN46, but that SARS-CoV-2 ORF7a localises additionally to small puncta (**Supplemental Figures 3A, 3B**). Golgi markers TGN46 and ZFPL1 were both found to be less reticular and fragmentation of Golgi markers was observed in HeLa cells stably expressing SARS-CoV-2 ORF7a (**Supplemental Figure 3C, 3D**). Such Golgi fragmentation is similarly observed in both HeLa and A549 cells infected with SARS- CoV-2 (**Figures 1E, 2E, 2F, Supplemental Figure 3E**)

To determine if ORF7a expression affected tetherin localisation, abundance, glycosylation or ability to form dimers, HeLa cells stably expressing SARS-CoV-1 ORF7a-FLAG or SARS-CoV2-ORF7a-FLAG were analysed by immunofluorescence, Western blotting and flow cytometry. By immunofluorescence ORF7a-FLAG and tetherin were not found to significantly overlap, but we did observe that the two proteins were both localised to some degree in adjacent perinuclear organelles (**Figure 4A**). Antagonism of tetherin by ORF7a could occur by interfering with tetherin’s ability to form homodimers. SARS-CoV-1 ORF7a was previously shown to alter glycosylation of ectopic tetherin without altering protein abundance, and the glycosylation-defective tetherin showed impaired ability to restrict SARS-CoV-1 egress [18]. To determine whether SARS-CoV-2 ORF7a impaired endogenous glycosylation or affected tetherin dimer formation, we performed Western blotting of WT HeLa, Bst2KO HeLa, KO + C3A-tetherin, and WT HeLa cells stably expressing either SARS-CoV-1 ORF7a-FLAG or SARS-CoV-2-ORF7a-FLAG (**Figure 4B**). C3A-tetherin stable cells (expressed in a tetherin KO HeLa background), are unable to form tetherin dimers and this can be observed by Western blotting. The stable expression of either SARS-CoV-1 ORF7a or SARS-CoV-2 ORF7a did not impact upon tetherin abundance, glycosylation, or the ability of tetherin to form dimers (**Figure 4B**). Finally, flow cytometry determined that expression of SARS-CoV-1 or SARS-CoV-2 ORF7a did not impact upon cell surface levels of tetherin (**Figure 4C**). Overall, these data show that ORF7a does not directly influence tetherin localisation, abundance, glycosylation or dimer formation.

**Figure 4.**
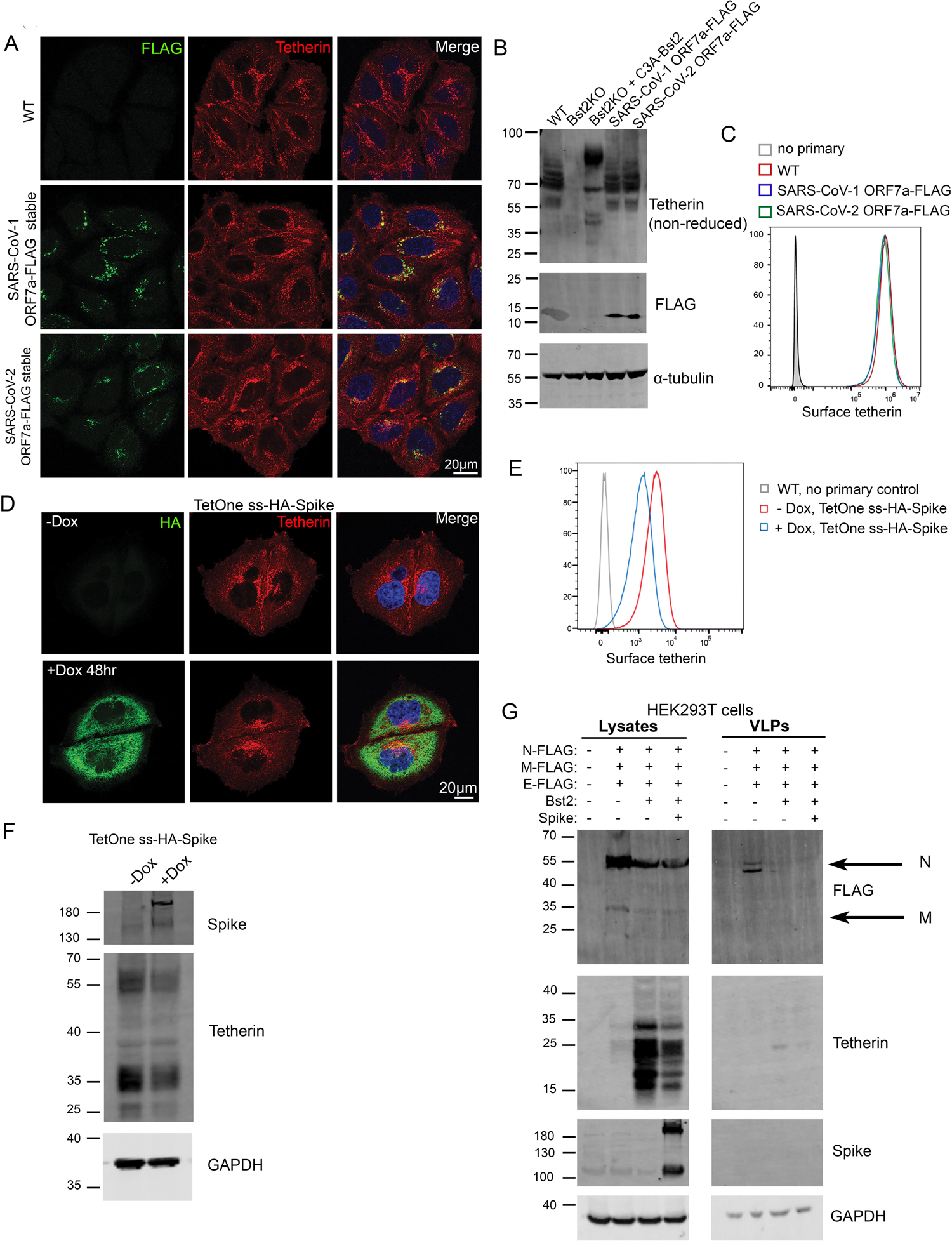
Spike but not ORF7a expression downregulates tetherin. (A) Stable SARS-CoV-1 ORF7a-FLAG and SARS-CoV-2 ORF7a-FLAG HeLa cell lines were generated and tetherin localization was analysed by immunofluorescence. Representative confocal immunofluorescence microscopy images of fixed wild-type, SARS-CoV-1 ORF7a-FLAG or SARS- CoV-2 ORF7a-FLAG HeLa cells were stained using anti-FLAG (green) and anti-tetherin (red) antibodies. (B) Western blotting was performed to confirm the abundance, glycosylation and dimer formation of tetherin upon stable expression of SARS-CoV-1 ORF7a- FLAG and SARS-CoV-2 ORF7a-FLAG. Bst2KO cells rescued with C3A-HA express tetherin that is unable to form homodimers. (C) Flow cytometry of wild-type (red), stable SARS-CoV-1 ORF7a-FLAG (blue) and stable SARS-CoV-2 ORF7a-FLAG (green) HeLa cells to analyse surface tetherin levels. (D) Stable tetracycline-inducible (TetOne) ss-HA-Spike HeLa cell lines were generated. Cells were incubated with Doxycycline for 48 hours to induce the expression of ss-HA-Spike. Representative confocal immunofluorescence image showing tetherin loss correlates with HA expression. Cells were stained using anti-HA (green) and anti-tetherin (red) antibodies. (E) Flow cytometry was performed on tetracycline-inducible ss-HA-Spike cells in resting conditions (no Doxcycline – red) or following induction (plus Doxycycline – blue). Surface tetherin levels were analysed. (F) Western blot analysis of tetherin in TetOne ss-HA-Spike HeLa cells in the absence of Doxycycline or following induction with Doxycycline for 48 hours. (G) Virus-like particle (VLP) experiments were performed using HEK293T cells. Cells were co-transfected with plasmids encoding N-FLAG, M-FLAG, E- FLAG, Bst2, Spike. Overall expression of FLAG was used to confirm transfection and VLP production. Whole cell lysates were collected and blotted, and VLPs were isolated from the culture supernatants. Blots were analysed using anti-FLAG, anti-tetherin, anti-Spike and anti-GAPDH (loading control) antibodies.

### Tetherin is downregulated by SARS-CoV-2 Spike

SARS-CoV-1 Spike causes tetherin downregulation via lysosomal degradation [19]. To interrogate the impact of SARS-CoV-2 Spike on tetherin, we generated an epitope tagged Spike construct to aid our imaging studies. We placed a HA epitope immediately after the native signal peptide sequence (ss-HA-Spike), rendering the HA tag at the N terminus of the mature protein. Transient transfection of cells with ss-HA-Spike caused a mild decrease in tetherin as observed by immunofluorescence (**Supplemental Figure 4A**), with tetherin being primarily lost from the plasma membrane. By flow cytometry, we observed no impairment in surface levels of ss-HA-Spike, or in the ability of ss-HA-Spike to bind ACE2 versus untagged Spike (**Supplemental Figure 4B, 4C**).

Attempts to generate stable cell lines constitutively expressing Spike were unsuccessful, most likely due to the accumulation of non-viable multinucleated cells (despite the lack of ACE2 expression), so we generated an inducible ss-HA-Spike cell line using the lentiviral TetOne system. Tetracycline-inducible ss-HA-Spike stable HeLa cells were generated, and expression of Spike was analysed by immunofluorescence microscopy following induction through Doxycycline treatment (**Figure 4D**). Doxycycline-induced ss-HA-Spike expression resulted in the rapid formation of numerous multi-nucleated syncytia (**Supplemental Figure 4D**), as reported by others [3,4,6]. Flow cytometry confirmed that Spike expression reduced the cell surface levels of tetherin (**Figure 4E**) and Western blotting revealed a mild downregulation of tetherin upon Spike expression (**Figure 4F**). To determine whether Spike specifically downregulates tetherin, or affects other proteins, we analysed the surface levels of another integral transmembrane protein, CD71, and found that the expression of Spike had a very minor impact on CD71 (**Supplemental Figure 4E**).

To explore whether the mild loss of tetherin induced by Spike expression would affect virus release, we performed experiments with virus like particles (VLPs) using HEK293T cells which do not express endogenous tetherin. VLPs were isolated from supernatants of cells expressing FLAG-tagged SARS-CoV-2 structural proteins (Nucleocapsid (N), Membrane (M), Envelope (E)). VLP yields were impaired by the additional expression of tetherin (**Figure 4G**). N-FLAG was most abundantly expressed in cells and recovered best in VLP fractions. The additional co-expression of Spike caused no change in VLP release, despite the reduction in cellular tetherin.

### ORF3a alters tetherin localisation by impairing retrograde traffic and enhances VLP release

Given the dramatic loss and relocalisation of tetherin in SARS-CoV-2 virally infected cells, yet only minor downregulation by Spike, we performed a miniscreen of SARS- CoV-2 ORFs to identify additional SARS-CoV-2 proteins involved in tetherin downregulation or antagonism. We transiently transfected HeLa cells with: ORF3a- Strep, ORF6-Strep, ORF7a-Strep, Strep-ORF7b, ORF8-Strep, ORF9b-Strep, Strep-ORF9c, ORF10-Strep and confirmed their expression by intracellular flow cytometry using an anti-Strep antibody (**Supplemental Figure 5A**). Surface tetherin levels were analysed by flow cytometry and no significant tetherin downregulation was observed upon expression of any ORF (**Figure 5A**). The intracellular tetherin localisation was analysed by confocal microscopy (**Supplemental Figure 5B**) and we noticed a redistribution of tetherin towards punctate organelles upon expression of ORF3a (**Figure 5B**).

**Figure 5.**
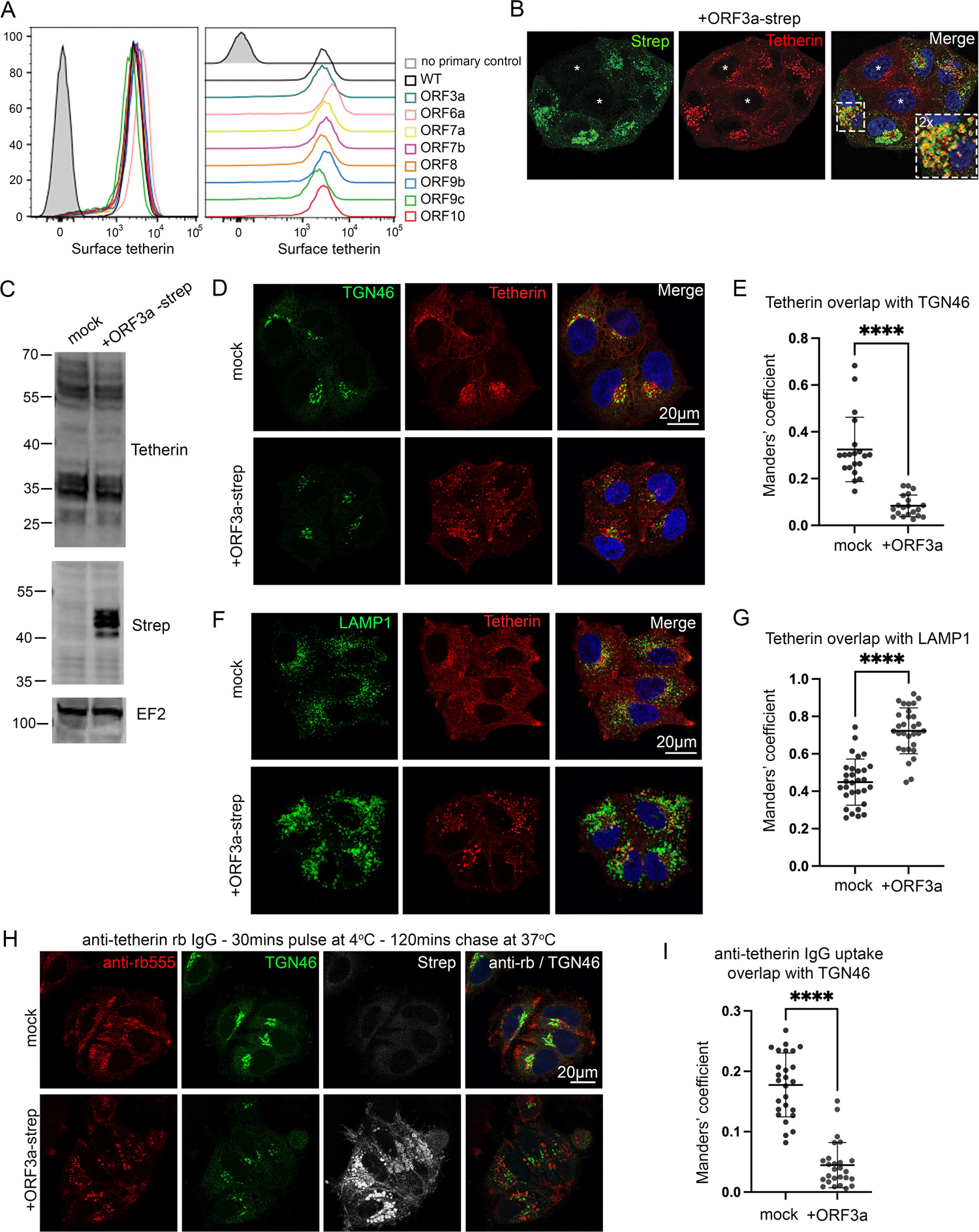
SARS-CoV-2 ORF3a redistributes tetherin away from the biosynthetic pathway via defective retrograde trafficking. (A) A miniscreen was performed to analyse the ability of other SARS-CoV-2 ORFs to downregulate surface tetherin. HeLa cells were transiently transfected with Strep-tagged plasmids encoding: ORF3a-Strep, ORF6-Strep, ORF7a-Strep, Strep-ORF7b, ORF8-Strep, ORF9b-Strep, Strep-ORF9c or ORF10-Strep. 48 hours post transfection, cells were stained for surface tetherin. (B) ORF3a-Strep transfected cells displayed intracellular tetherin accumulation. Tetherin accumulated in ORF3a-Strep positive compartments. Representative confocal immunofluorescence microscopy image of fixed HeLa cells transiently transfected with SARS-CoV-2 ORF3a-Strep. Cells were fixed and stained with anti-Strep (green) and anti-tetherin (red) antibodies and DAPI (blue). (C) HeLa cells were transiently transfected with SARS-CoV-2 ORF3a-Strep. Mock and transfected cells were lysed 48 hours post transfection and analysed by Western blot. Blots were analysed using anti-tetherin, anti-Strep and anti-EF2 (loading control) antibodies. (D) Confocal immunofluorescence microscopy was performed on mock or ORF3a-Strep transiently transfected HeLa cells to analyse tetherin overlap with TGN46. Cells were fixed and stained using anti-TGN46 (green), anti- tetherin (red) antibodies, and DAPI (blue). Representative images are shown. (E) Colocalisation analysis was performed to quantify the Mander’s overlap coefficient of tetherin overlapping TGN46. At least 20 cells per condition from three independent experiments were analysed. Individual data points are plotted with mean and standard deviation. Two-tailed, unpaired t-tests were performed. **** p < 0.0001. (F) Confocal immunofluorescence microscopy was performed on mock or ORF3a-Strep transiently transfected HeLa cells to analyse tetherin overlap with the lysosomal marker, LAMP1. Cells were fixed and stained using anti-LAMP1 (green), anti-tetherin (red) antibodies, and DAPI (blue). Representative images are shown. (G)Colocalisation analysis was performed to quantify the Mander’s overlap coefficient of tetherin overlapping LAMP1. At least 30 cells per condition from three independent experiments were analysed. Individual data points are plotted with mean and standard deviation. Two-tailed, unpaired t-tests were performed. **** p < 0.0001. (H) Antibody uptake experiments were performed to investigate the fate of endocytosed tetherin. Transient transfections were performed 48 hours prior to uptake experiments. Anti-tetherin antibodies were bound to live cells on ice for 30 minutes before a 2 hour chase at 37°C. Cells were fixed and immunolabelled using anti-TGN46 (green), anti-Strep (white), secondary anti- rabbit555 (red) antibodies, and DAPI (blue). Representative images are shown. (I) Colocalisation analysis was performed to quantify the Mander’s overlap coefficient of endocytosed anti-tetherin overlapping TGN46. At least 26 cells per condition from three independent experiments were analysed. Individual data points are plotted with mean and standard deviation. Two- tailed, unpaired t-tests were performed. **** p < 0.0001.

SARS-CoV-2 ORF3a is a viroporin that localises to and damages endosomes and lysosomes [29]. As such, the observed increased presence of tetherin puncta following ORF3a expression may be due to decreased lysosomal degradation. Western blotting showed that expression of OFR3a had no impact on total levels of endogenous tetherin protein (**Figure 5C**). Flow cytometry confirmed minimal differences in tetherin levels upon transient expression of ORF3a (**Supplemental Figure 6A**). We noticed that tetherin localisation appeared altered upon ORF3a expression and observed a decrease in perinuclear tetherin that appeared in and around the Golgi (**Figure 5D**), and that tetherin localised to ORF3a-Strep positive punctate organelles. We quantified the levels of colocalisation between tetherin and the trans-Golgi marker, TGN46, and found a significant loss of tetherin from this region upon expression of ORF3a (**Figure 5E**). The loss of tetherin from the peri- Golgi area in ORF3a transfected cells was associated with an increase in tetherin within LAMP1 positive punctate organelles (**Figure 5F, 5G**). Expression of ORF3a also disrupted the distribution of numerous endosome-related markers including CIMPR, VPS35, CD63, (**Supplemental Figure 6B**), and the mixing of early and late endosomal markers EEA1 and cathepsin D (**Supplemental Figure 6C**). By TEM, ORF3a expressing cells contained enlarged endolysosomes, consistent with defects in endolysosomal homeostasis (**Supplemental Figure 6D**).

The ORF3a-mediated increase in tetherin abundance within lysosomes could be due to defective lysosomal degradation but could also be due to altered intracellular trafficking. In addition to the plasma membrane, tetherin is found in several intracellular organelles, including ERGIC, Golgi and endosomes. The persistence of proteins within biosynthetic organelles at steady state can be due to the presence of retention motifs or retrograde trafficking [30] – where proteins are recovered from endosomes back to the biosynthetic pathway.

Tetherin has recently been shown to undergo some retromer-dependent retrograde trafficking [31]. We investigated whether ORF3a could impair the retrograde traffic of tetherin. Antibody uptake experiments were performed and the localisation of the internalised antibody assessed by immunofluorescence. In mock cells, fed anti- tetherin antibodies were found at the cell periphery and in the peri-Golgi region (**Figure 5H**), confirming retrograde recycling of tetherin. However, in cells expressing ORF3a, endocytosed tetherin was not recovered to the peri-Golgi region (**Figure 5H).** Colocalisation analysis confirmed a significant loss of anti-tetherin labelling overlapping with TGN46 upon the expression of ORF3a (**Figure 5I**). Internalised anti-tetherin antibodies were clearly found to colocalise with LAMP1 in ORF3a-Strep transfected cells (**Supplemental Figure 6E**), demonstrating that tetherin recycling was inhibited.

To establish whether the loss of retrograde traffic of tetherin was cargo specific, we similarly performed antibody pulse-chase experiments using the well-established retrograde cargo, CIMPR (**Supplemental Figures 6F, 6G**). Similar phenotypes were observed, indicating that ORF3a-Strep expression impairs global retrograde recycling and is not specific to tetherin.

**Figure 6.**
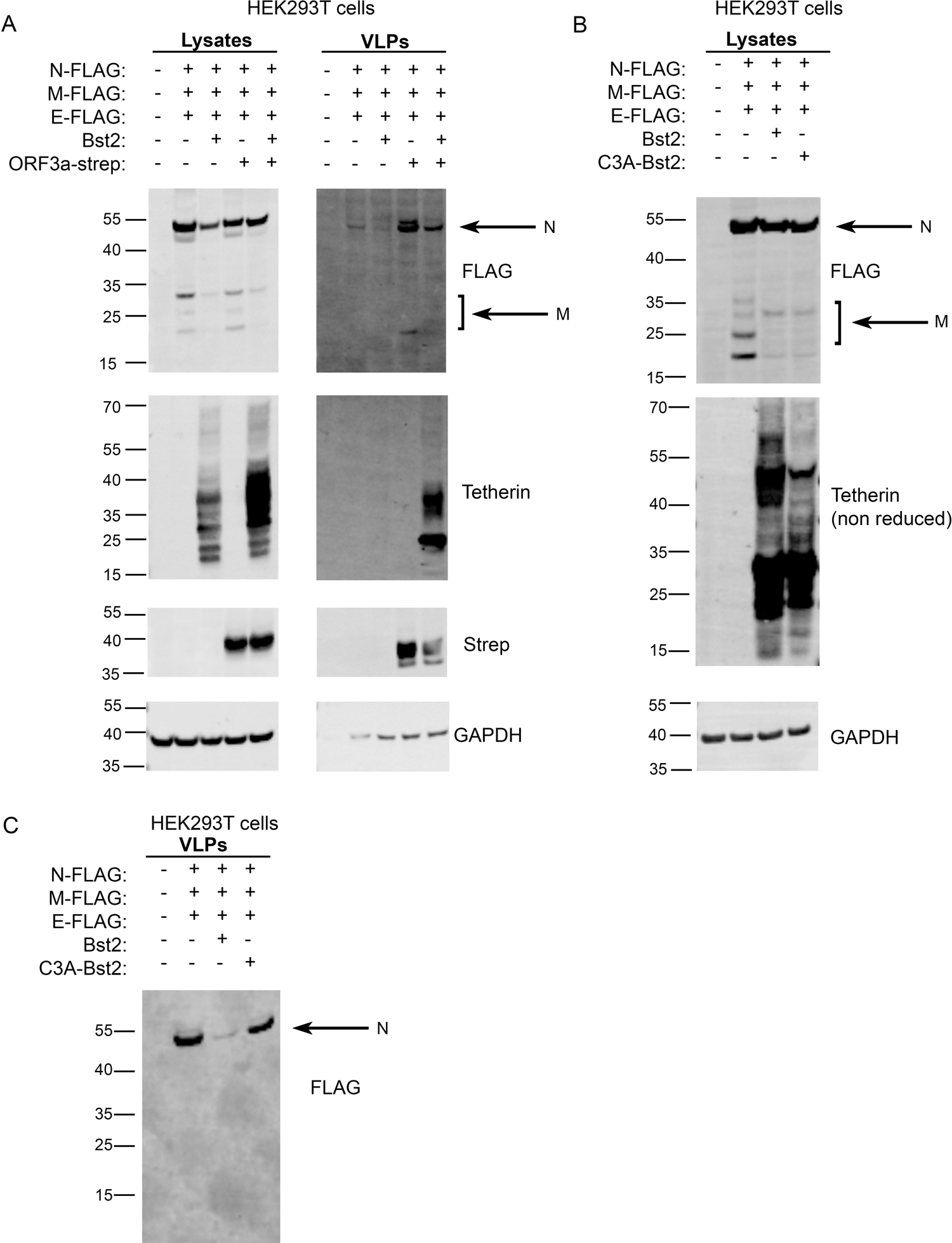
SARS-CoV-2 ORF3a enhances VLP release despite enhancing cellular tetherin. (A) Virus-like particle (VLP) experiments were performed using HEK293T cells. Cells were co-transfected with plasmids encoding N-FLAG, M-FLAG, E- FLAG, Bst2, ORF3a-Strep. Overall expression of FLAG was used to confirm structural protein transfection and VLP production. Whole cell lysates were collected, and VLPs were isolated from the culture supernatants. Blots were analysed using anti-FLAG, anti-tetherin, anti-Strep and anti-GAPDH (loading control) antibodies. (B) Experiments were performed to analyse the effect of tetherin-dependent reuptake and degradation of structural proteins. Cells were co-transfected with plasmids encoding N-FLAG, M-FLAG, E-FLAG, Bst2, C3A-Bst2. Whole cell lysates were collected. Blots were analysed using anti-FLAG, anti-tetherin and anti- GAPDH (loading control) antibodies. (C) VLPs were collected from (B) and the levels of VLP release was analysed using anti- FLAG antibodies.

To determine whether the ORF3a-mediated loss of tetherin from biosynthetic organelles altered SARS-CoV-2 egress, we again performed VLP experiments in HEK293T cells. VLPs were detected from the supernatants of cells expressing N, M and E and yields were significantly reduced upon addition of tetherin (**Figure 6A**).

The amount of VLPs were greatly increased by the expression of ORF3a alone, indicating that ORF3a may enhance VLP release independently of its antagonism of tetherin. The co-expression of structural proteins, tetherin and ORF3a resulted in a lower yield of VLP when compared with structural proteins and ORF3a only.

We reproducibly found that the expression of tetherin reduced cellular levels of structural proteins (**Figures 4G, 6A**), and we hypothesised that tetherin expression could promote endocytosis and subsequent degradation of VLPs. Tetherin restriction of enveloped viruses prevents their egress, but also promotes internalisation of tethered virions which are redirected to lysosomal degradation. To test this, we expressed either wildtype tetherin or C3A-tetherin which is unable to form homodimers and therefore unable to promote viral retention and reinternalisation. Both WT and C3A-tetherin gave similar reductions in cellular levels of structural proteins (**Figure 6B**), indicating that reduced structural protein levels were independent of tethering, reinternalization and subsequent degradation. As expected, the yield of VLPs from C3A-tetherin expressing cells was similar to that when no tetherin was expressed, confirming that tetherin homodimers are required for SARS-CoV-2 restriction (**Figure 6C**).

## Discussion

Tetherin retains the ability to restrict a number of different enveloped viruses that bud at distinct organelles. Tetherin forms homodimers that sit on opposing membranes (e.g., plasma membrane and viral envelope) that are linked via disulphide bonds formed between three luminal cysteine residues. For tetherin to restrict SARS-CoV- 2, tetherin molecules must be incorporated into the virus during the process of envelopment, which occurs via a single step in modified ERGIC organelles. The removal of tetherin from the respective budding compartments of different enveloped viruses appears as a central feature of tetherin antagonism. For example, during HIV-1 infection tetherin is downregulated from the plasma membrane – the budding compartment for HIV-1 – and is retained in the TGN by the action of HIV-1 Vpu [32]. The opposing relocalisation of tetherin by SARS-CoV-2 – *away* from the biosynthetic organelles – similarly reflects the need of SARS-CoV-2 to remove tetherin from its budding compartment.

Our data support that tetherin molecules become incorporated to SARS-CoV-2 virions and act to restrict virus release upon delivery of virions to the plasma membrane. SARS-CoV-2 counteracts this restriction in a variety of ways, ensuring its continued transmission. We have also shown that ORF3a antagonises tetherin by altering its steady state distribution, and that VLP yields are enhanced upon ORF3a expression (**Figure 6A**). We also demonstrate that Spike downregulates tetherin, consistent with reports for SARS-CoV-1 [19], but we found this downregulation to be insufficient to promote VLP release (**Figure 4G**).

Tetherin is an IFN-stimulated gene (ISG) [13], and many cell types express low levels of tetherin at resting states. Weak type I IFN responses appear as a hallmark of sarbecovirus infections - SARS-CoV-1 is a poor inducer of type I IFN [33], and SARS-CoV-2 weaker still. By using cell lines that ubiquitously express high levels of tetherin and cell lines that require induction with IFN, we have demonstrated that SARS-CoV-2-mediated tetherin downregulation is not solely dependent upon dysregulation of IFN responses, and that other mechanisms exist to antagonise tetherin.

SARS-CoV-2 infection causes dramatic rearrangements of intracellular organelles (**Figures 1E, 2D, 2E, 2F**), the most striking of which is the rearrangement of the late biosynthetic compartments and formation of DMVs where viral particles are assembled. In addition to the clear perturbation to the biosynthetic organelles we also observed dysfunction within endolysosomal organelles. Expression of SARS- CoV-ORF3a impaired retrograde traffic, reducing the amount of tetherin retrieved to biosynthetic organelles, and permitting VLP release (**Figures 5D-I, 6A**). ORF3a dimers form non-selective calcium permeable cation channels [34] that localise to endocytic organelles [35]. These pores disrupt endolysosomal acidification [35] and promote lysosome fusion with the plasma membrane [36], which may aid SARS- CoV-2 lysosomal egress [35]. Secretion of lysosomal hydrolases has been observed upon ORF3a expression [31] and whilst this may in-part be due to enhanced lysosome-plasma membrane fusion, our finding that ORF3a impairs CIMPR retrieval may also be a contributing factor (**Supplemental Figures 6B, 6F, 6G**).

Precisely how ORF3a impedes retrograde traffic is unclear. Recruitment of VPS35, a core retromer component, to membranes does not appear impeded by ORF3a expression (**Supplemental Figure 6B**), indicating that tubule formation or scission may be impaired. Endosomal acidification has been proposed to regulate tubule formation, as a homeostatic mechanism to ensure tubulation doesn’t occur too early in the endocytic pathway or from incorrect organelles [37].

The ORF3a-mediated defective retrograde trafficking is not specific to tetherin, and likely affects numerous other protein cargos, some of which may aid the infectivity and transmissibility of SARS-CoV-2. Our findings highlight defective retrograde traffic as a novel mechanism of tetherin antagonism. Whether other enveloped viruses which bud within the biosynthetic pathway employ similar strategies remains to be investigated.

In addition to our identification of the role of ORF3a in tetherin antagonism, we also investigated the roles of two previously identified tetherin antagonists – ORF7a and Spike. We did not observe any difference in total tetherin levels, tetherin glycosylation, ability to form dimers, or surface tetherin upon expression of SARS- CoV-2 ORF7a (**Figures 4A, 4B, 4C**). The intracellular localisation of tetherin also appeared unaffected by ORF7a expression. However, others have demonstrated a role for ORF7a in sarbecovirus infection and both SARS-CoV-1 and SARS-CoV-2 virus lacking ORF7a show impairments in virus replication in the presence of tetherin [18, 38]. A direct interaction between SARS-CoV-1 ORF7a and SARS-CoV-2 ORF7a and tetherin have been described [18, 38], although the precise mechanism(s) by which ORF7a antagonises tetherin remains enigmatic, as does its role in pathogenesis. A number of SARS-CoV-2 variants have been described which contain truncated ORF7a, including the so-called ‘Arizona’ ORF7a [39] – lacking a signal sequence, and ORF7a^Δ115^ [40] which contains a frameshift mutation leading to a premature stop codon, preventing translation of the transmembrane domain and cytosolic tail.

SARS-CoV-2 Spike has well documented roles in facilitating SARS-CoV-2 viral entry, and in syncytia formation [41]. We found that SARS-CoV-2 Spike expression caused a mild downregulation of tetherin from cells, consistent with reports from SARS-CoV-1 [19], although in our experiments the downregulation was not sufficient to promote VLP release (**Figure 4G**). Overexpression of SARS-CoV-2 Spike alone can induce the unfolded protein response (UPR) [42], and this may be responsible for the mild downregulation of tetherin.

SARS-CoV-2 spreads through cell-to-cell transmission in a process mediated by Spike [43]. This cell-to-cell transmission is sensitive to endosomal entry inhibitors, implicating endosomal membrane fusion as a mechanism for infection. Tetherin impairs the egress of viruses from cells, but for many viruses also impairs cell-to-cell transmission. HIV-1 viral aggregates are not efficiently endocytosed by cells and their fusion capacities reduced [25,44,45].

The multiple mechanisms by which tetherin is antagonised by different enveloped viruses highlights the importance of overcoming restriction in determining the success of enveloped viruses.

## Materials and methods

### Antibodies

Primary antibodies used in the study were:

rat anti-DYKDDDDK (L5) (BioLegend, WB 1:1000, IF 1:200); rat anti-HA (Roche, 3F10, IF 1:500); rabbit monoclonal anti-tetherin (Abcam, ab243230, WB 1:2000, IF 1:400, surface EM 1:200); mouse anti-SARS-CoV-2 Spike antibody 1A9 (GeneTex, GTX632604, WB 1:1000, IF 1:300); rabbit anti-TGN46 (Abcam, ab50595, 1:300); rabbit anti-ZFPL1 (Sigma-Aldrich, HPA014909, 1:500); rabbit anti- Beta2microglobulin (Dako 1:500); mouse anti-ACE2 (Proteintech, 66699, WB 1:1000), Phycoerythrin-conjugated anti-human tetherin (BioLegend, RS38E, FC 2 μg / 100 μl / 10^6^ cells), Pacific Blue-conjugated anti-CD71 (Exbio, MEM-75, FC 1 μg / 100 μl / 10^6^ cells), StrepMAB-Classic (IBA LifeSciences, WB 1:2000, IF 1:500), StrepMAB-Classic DY-549 (IBA LifeSciences, FC 1:500), Strep-Tactin-DY649 (IBA LifeSciences, IF 1:2000), mouse anti-CIMPR (2G11) (Abcam, ab2733, IF 1:200), mouse anti-VPS35 (Santa Cruz, B-5, IF 1:300), mouse anti-CD63 (BioLegend, H5C6, IF 1:300), mouse anti-EEA1 (BD Transduction, 610457, IF 1:200), rabbit anti- Cathepsin D (Calbiochem, 219361, IF 1:500), rabbit anti-GAPDH (Cell Signalling 14C10, WB 1:2000), goat anti-EF2 (Santa Cruz, C-14, WB 1:10,000), mouse anti- tubulin (Proteintech, 66031, WB 1:10,000)

Secondary antibodies used in this study were:

Goat anti-Mouse IgG Alexa488/555 and Goat anti-rabbit IgG Alexa 488/555 (ThermoFisher) secondary antibodies were used for confocal microscopy.

Goat IRDye 680 anti-mouse, anti-rabbit, anti-goat, anti-rat and Goat IRDye 800 anti- mouse, anti-rabbit antibodies (Li-Cor) were used for Western blotting.

### Cloning

pcDNA6B SARS-CoV-1 and SARS-CoV-2 ORF7a-FLAG constructs were a gift from Professor Peihui Wang (Shandong University, China). To generate stable cell lines, ORF7a-FLAG cDNA fragments were subcloned into pQCXIH retroviral vectors. ss-HA-Spike was generated by cloning an HA epitope plus a Serine-Glycine linker between residues S13 and Q14 of SARS-CoV-2 spike. Following translocation to the ER lumen, cleavage of the signal sequence will render the HA tag at the N terminus of the mature protein. Codon optimised SARS-CoV-2 spike was a gift from Dr Jerome Cattin/Professor Sean Munro (LMB, Cambridge, UK). ss-HA-Spike was originally cloned to pcDNA6B for transient transfection, and subsequently subcloned into pLVX-TetOne. Gene expression from TetOne stable cell lines was induced by treatment with Doxycycline (1 μg/mL final concentration – Caymen Chemical) All cloning was verified by Sanger sequencing (GeneWiz).

The soluble ACE2 plasmid (pcDNA3-sACE2(WT)-Fc(IgG1) was a gift from Dr Erik Procko (Addgene plasmid: 145163) [46] The StrepTactin tagged SARS-CoV-2 M protein plasmid (pLVX-EF1alpha-nCoV2019-M-IRES-Puro), and StrepTactin tagged SARS-CoV-2 ORF library was a gift from Dr David Gordon [47].

Bst2-C3A-HA was a gift from Prof George Banting (University of Bristol) and the cDNA was cloned to pQCXIH for stable expression in Bst2KO HeLa cells.

### Immortalised cell lines

A549 cells were a gift from Dr Brian Ferguson, University of Cambridge, UK and were cultured in DMEM supplemented with 10% fetal bovine serum, L-glutamine, and Penicillin/Streptomycin, in 5% CO2 at 37 °C. T84 cells were purchased from ATCC and were cultured in DMEM: F-12 medium containing 5% fetal bovine serum, L-glutamine, and Penicillin/Streptomycin, in 5% CO2 at 37 °C. HeLa cells were a gift from Professor Scottie Robinson (CIMR, University of Cambridge, UK) and were cultured in DMEM supplemented with 10% fetal bovine serum, L-glutamine, and Penicillin/Streptomycin, in 5% CO2 at 37 °C. Bst2KO HeLa cells were previously described [27]. All cells were tested for mycoplasma tested using MycoAlert Mycoplasma Detection Kit (Lonza).

### Human airway epithelium (HAE) cells

Nasal brushing samples were taken from healthy participants turbinate using a 3-mm bronchial cytology brush under Health Research Authority study approval (REC ref: 20/SC/0208; IRAS: 282739). The nasal brushings were placed in PneumaCult-Ex Plus medium (STEMCELL Technologies, Cambridge, UK) and cells extracted from the brush by gentle agitation. The cells were seeded into a single well of a collagen (PureCol from Sigma Aldrich) coated plate and once confluent, they were passaged and expanded further in a T25 flask. The cells were passaged a second time and seeded onto Transwell inserts (6.5 mm diameter, 0.4 μm pore size, Corning) at a density of 24,000 cells per insert. Cells were cultured in PneumaCult-Ex Plus medium (STEMCELL Technologies, Cambridge, UK) until confluent before replacing with PneumaCult-ALI medium in the basal chamber and the apical surface exposed, giving an air liquid interface to stimulate cilia biogenesis.

### SARS-CoV-2 infections of HAE cells

SARS-CoV-2 isolate hCoV-19/England/204501206/2020 (EPI_ISL_660791) was isolated from swabs as described in [48]. The isolate was passaged twice in Vero cells before being used to infect HAE cells. To remove the mucus layer from the apical surface of HAE cells prior to infection, 200 μL of DMEM was added to the HAE cells at 37 °C, 5% CO2 for 10 minutes. The cells were infected at a MOI of 0.01 with inocula added to the apical chamber and incubated for 1 hour at 37 °C, 5% CO2 before removal of the inoculum and incubating for a further 48 hours.

### Transient transfections

HeLa cells were transfected with 2.5 μg of DNA using TransIT-HeLa Monster (Mirus Bio) according to the manufacturer’s instructions. Cells were analysed 48 hours after transfection.

### SARS-CoV-2 ORF miniscreen

HeLa cells were transiently transfected with pLVX-EF1alpha-nCoV2019 2xStrep- tagged SARS-CoV2 ORF constructs [47] using TransIT-HeLa Monster (Mirus Bio). After 48 hours transfection, cells were detached and the cell population equally split. Half of the cells were permeabilised and intracellular Strep levels were measured by flow cytometry. Surface tetherin staining was performed on the remaining half of the cells.

### Generation of stable cell lines

#### Lentiviral constructs

ACE2 stable HeLa and A549 cell lines were generated using the lentiviral pLVX- ACE2-Blasticidin construct from Dr Yohei Yamauchi (University of Bristol). Following transduction, cells were selected with 10 μg/mL blasticidin for 18 days.

Expression-inducible stable cell lines were generated using the pLVX-TetOne system. pLVX-TetOne-Puro-ORF7a-2xStrep and pLVX-TetOne-Puro-2xStrep- ORF9c were a gift from Dr David Gordon (UCSF, USA). pLVX-TetOne-Puro-ss-HA- spike was generated as described above. Following transduction, cells underwent antibiotic selected with 1 μg/mL puromycin for 5 days.

HEK293T cells were transfected with lentiviral vectors (pLVX / pLVX-TetOne) plus packaging plasmids pCMVR8.91 and pMD.VSVG using TransIT-293 (Mirus Bio). Viral supernatants were collected 48 hours after transfection, passed through 0.45 μm filters and recipient cells transduced by ‘spinfection’ – viral supernatants were centrifuged at 1800 rpm in a benchtop centrifuge at 37 °C for 3 hours to enhance viral transduction.

#### Retroviral constructs

pQCXIH-SARS-CoV-1-ORF7a-FLAG and pQCXIH-SARS-CoV-2-ORF7a-FLAG were transfected to HEK293T cells with the packaging plasmids pMD.GagPol and pMD.VSVG using Trans-IT293 (Mirus Bio). Viral supernatants were collected 48 hours after transfection, passed through 0.45 μm filters and recipient cells transduced by ‘spinfection’ – viral supernatants were centrifuged at 1800 rpm in a benchtop centrifuge at 37 °C for 3 hours to enhance viral transduction. Stable cells were selected using 400 μg/mL Hygromycin B for 10 days.

### SARS-CoV-2 infections of immortalised cell lines

WT HeLa+ACE2, Bst2KO HeLa+ACE2, A549+ACE2 or T84 cells were infected with isolate BetaCoV/ Australia/VIC01/2020 [22], which had been passaged once on Vero cells following receipt from Public Health England. All cells were washed with PBS before being infected with a single virus stock, diluted to the desired MOI with sera- free DMEM (supplemented with 25mM HEPES, penicillin (100 U/mL), streptomycin (100 g/mL), 2 mM L-glutamine, 1 % non-essential amino acids). After one hour, the inoculum was removed, and cells washed again with PBS. Infected cells were maintained in DMEM, supplemented with the above-described additions plus 2% FCS (virus growth media).

For immunofluorescence, cells were plated to glass-bottomed 24-well plates (E0030741021, Eppendorf) and infected at an MOI of 0.5 and incubated for 24 hours, after which plates were submerged in 4% PFA/PBS for 20 min.

For conventional electron microscopy, cells were plated to plastic Thermanox (Nunc) coverslips in 24-well plates and infected at an MOI of 0.5 and incubated for 24 hours, after which plates were submerged in 2% PFA / 2.5% glutaraldehyde / 0.1M cacodylate buffer for 20 minutes.

For surface labelling immunoEM, cells were plated to Thermanox coverslips, infected at an MOI of 0.5 and incubated for 24 hours, and fixed with 4% PFA / 0.1M cacodylate buffer for 20 minutes.

### Conventional electron microscopy

Cells were fixed (described above) before being washed with 0.1M cacodylate buffer. Cells were stained using 1% osmium tetroxide + 1.5% potassium ferrocyanide fo r 1 hour before staining was enhanced with 1% tannic acid / 0.1M cacodylate buffer for 45 minutes. Cells were washed, dehydrated and infiltrated with Epoxy propane (CY212 Epoxy resin:propylene oxide) before being embedded in Epoxy resin. Epoxy was polymerised at 65 °C overnight before Thermanox coverslips were removed using a heat-block. 70nm sections were cut using a Diatome diamond knife mounted to an ultramicrotome. Ultrathin sections were stained with UA Zero (Agar scientific) and lead citrate. An FEI Tecnai transmission electron microscope at an operating voltage of 80kV was used to visualise samples, mounted with a Soft Imaging System Megaview III digital camera.

### Surface immunogold labelling

To enable luminal surface epitopes to be labelled, cells were fixed with 4% PFA / 0.1 M cacodylate. They were washed with 0.1 M cacodylate buffer before being blocked with 1% BSA/PBS. Coverslips were inverted over drops of either rabbit anti-tetherin or rabbit anti-spike antibodies diluted in 1% BSA/PBS. Coverslips were washed before being incubated with protein A gold. Following gold labelling, cells were re- fixed using 2% PFA / 2.5% glutaraldehyde / 0.1 M cacodylate before being processed for conventional electron microscopy as described above.

### Immunofluorescence microscopy

A549, T84 or HeLa cells were grown on glass coverslips and fixed using 4% PFA/PBS. Cells were quenched with 15 mM glycine/PBS and permeabilised with 0.1% saponin/PBS. Blocking and subsequent steps were performed with 1% BSA, 0.01% saponin in PBS. Cells were mounted on slides with mounting medium containing DAPI (Invitrogen). Cells were imaged using a LSM700 confocal microscope (63×/1.4 NA oil immersion objective; ZEISS).

### Immunofluorescence of cryostat sections

HAE cells were fixed in 4% PFA / PBS for 1 hour before embedding and freezing in OCT (optimal cutting temperature) compound. ∼15 μm thick sections were cut using a cryostat and these permeabilised using 0.2% saponin in PBS for 20 minutes at room temperature before incubating in blocking solution containing 0.02%, 1% BSA in PBS for 30 minutes at room temperature. The cryostat sections were incubated with primary antibodies in blocking solution for 2 hours at room temperature.

Subsequently, Alexa Fluor bound secondary antibodies and phalloidin 647 (Abcam, ab176759) were applied to the sections for 1 hour at room temperature. Images were acquired using a Leica SP8 confocal.

### Immunofluorescence colocalisation analysis

Appropriate threshold values were manually applied to each channel and the Manders’ overlap coefficient between two channels was quantified using the JACoP plugin of ImageJ Fiji software [49]. The same threshold values were applied to all the cells quantified. For analysis of ORF3a transient transfected cells, transfected cells were identified by anti-Strep-647 labelling and cell masks were manually drawn and applied to each channel for analysis.

### Antibody uptake experiments

Cells were seeded to glass coverslips in 24-well plates a day before uptake experiments were performed. Antibody was diluted in pre-chilled 1% BSA / PBS (to 5 μg/mL) and 50μL drops pipetted to parafilm-covered blocks on ice. Coverslips were removed from 24-well plates are placed cell-side down to diluted antibody drops on ice. Cells were incubated on ice for 30 mins to allow antibodies to bind to proteins at the cell surface. Coverslips were then removed and placed back in 24-well plates, and transferred back to a 37°C, 5% CO2 incubator to allow antibody-antigen complexes to be endocytosed and trafficked for 2 hours. Coverslips were then fixed and processed for immunofluorescence microscopy using secondary antibodies specific for internalised antibodies to detect antibody localisation.

### Western blotting

Tetherin blots were performed using Laemmli sample buffer and run in non-reducing conditions as previously described [25]. For all other blots, lysates were mixed with 4x NuPage LDS sample buffer (ThermoFisher). Gels were loaded to NuPage 4-12% Bis-Tris precast gels (ThermoFisher) and transferred to PVDF membranes before being blocked using 5% milk / PBS / 0.1% Tween. Primary antibodies and secondary antibodies were diluted in PBS-tween. Blots were imaged using an Odyseey CLx (Li- Cor).

### Virus Growth Curves

Multiple subconfluent T25 flasks of WT HeLa+ACE2 and Bst2KO HeLa+ACE2 cells were each infected with a single stock of SARS-CoV-2 (BetaCoV/Australia/VIC01/2020), at an MOI of 5 and 1, for the one-step and multi- step growth curves, respectively. After one hour of infection, cells were washed with PBS and maintained in 5 mL virus growth media until harvest. One flask was harvested at each time point (0-, 24-, 48- and 72-hours post-infection). At each time point, the supernatant (containing released virions) was collected, clarified and stored at −80 °C. The cell monolayer was scraped into 2 mL PBS and subjected to three freeze-thaw cycles to release intracellular virions. Following clarification, the cell debris was discarded, and the remaining supernatant was stored at −80 °C. The infectious titre of all virus samples was determined by plaque assay.

### Plaque assays

Plaque assays were performed as previously described for SARS-CoV-1, with minor amendments [50, 51]. Briefly, subconfluent Vero cells in 6-well plates were infected with serial dilutions of the virus sample, diluted in sera-free media, for one hour with constant rocking. After removal of the inocula and washing with PBS, 3 mL of 0.2% agarose in virus growth media was overlaid and the cells were incubated at 37 °C for 72 hours. At this time the overlay media was removed, cells were washed with PBS and fixed with 10% formalin, before being stained with toluidine blue. Plaques were counted manually.

### Flow cytometry

Cells were gently trypsinised, and surface stained for flow cytometry in PBS with 0.5 % BSA + 1 mM EDTA (FACS buffer) for 30 min on ice. For intracellular staining, surface stained cells were fixed and stained with Foxp3 / Transcription Factor Staining Buffer Set (eBioscience) according to manufacturer’s protocol. Samples were acquired on a four laser Cytoflex S (Beckman coulter, 488 nm, 640 nm, 561 nm, 405 nm).

### Flow Cytometry method to assess ACE2 binding

To generate soluble ACE2-Fc, approximately 5 million HEK-293T cells were transfected with 10 μg of pcDNA3-sACE2(WT)-Fc(IgG1) complexed to 50 μg of PEI. 72 hours post transfection the tissue culture supernatant was collected from the cells and any debris removed via centrifugation. The media was then aliquoted and frozen at -20 °C.

To determine the cell surface levels of the recombinant spike constructs, approximately 200,000 HEK-293T cells were transfected with 1 μg of plasmid DNA complexed to 3 μL of FuGENE HD (Promega, Cat: E2311). 16-20 hours post transfection the cells were dissociated from the plastic using cell dissociation buffer (Gibco Cat: 13151014) and resuspend in complete media. The cells were incubated with a SARS-CoV-2 spike antibody (GeneTex Cat: GTX632604; 1:400 diluted in media) for 1 hour on ice. Unbound antibody was removed by washing the cells with ice cold media and the bound antibody detected using a goat anti-mouse secondary antibody conjugated to Cy5 (Jakson ImmunoResearch Cat:115-175-146; diluted 1:500 in media) for 1 hour on ice. The cells were then washed three times with ice cold media and their fluorescent intensity measured using a BD FACSCalibur flow cytometer. Live cells were gated using forward/side scatter and approximately 10,000 events collected per sample.

To measure the ability of the recombinant spike constructs to bind ACE2, transfected cells (see previous section) were incubated with tissue culture supernatant containing ACE2-Fc for 1 hour on ice. Unbound ACE2-Fc was removed by washing the cells with ice cold media and the bound ACE2 detected using goat anti-human secondary antibody conjugated to Alexa647 (ThermoFisher Scientific Cat: A21445; diluted in media at 1:500) for 30 minutes on ice. The cells were then washed three times with ice cold media and their fluorescent intensity measured as outlined previously. To determine the specificity of the ACE2-Fc binding non-transfected cells were also incubated with the ACE2-Fc and the anti-human secondary antibody.

### VLP assays

HEK293T cells were seeded to 9 cm dishes. The plasmid constructs were co- transfected with TransIT-293 (Mirus Bio) and 5 μg of each plasmid (pcDNA3.1 N- FLAG, pcDNA3.1 M-FLAG, pcDNA3.1 E-FLAG, pQCXIH Bst2-HA, pQCXIH Bst2-C3A-HA, pcDNA3.1 Spike (codon optimised), pLVX ORF3a-Strep). SARS-CoV-2 VLPs were harvested 48 hours post transfection. VLPs were purified from culture supernatants by centrifugation at 500 x*g*, 10 mins, followed by a second centrifugation at 2,000 x*g* for 20 mins. Collected supernatants were filtered through 0.45 um filters before filtrates were layered on top of 20% sucrose cushions, and then centrifuged at 100,000 x*g* for 3 hours. The final pellets were resuspended in pre-chilled lysis buffer.

## Acknowledgments

We wish to thank the microscopy and flow cytometry facilities at the Department of Pathology, University of Cambridge and the electron microscopy facility at Cambridge Institute for Medical Research, University of Cambridge. We also thank Ranjit K. Rai and Paul Griffin for assistance with HAE cell culture and Anand Shah for help to gain ethical approval to acquire HAE cells for this study. Viral infection of HAE cells was done in the Wendy S. Barclay lab (Imperial College London) supported by G2P-UK National Virology Consortium funded by UKRI (to Jonathan C. Brown and Wendy S. Barclay). We wish to thank Melbourne Health and Public Health England for nCoV/Victoria/1/2020 Coronavirus strain.

## Author contributions

JRE conceived the study. JRE performed IF, EM, molecular and biochemical experiments. HS performed all work with SARS-CoV-2 infections, including viral growth curves and plaque assays with help from GWC. TB and JB performed HAE infection experiments. Flow cytometry was performed by KHJ, NMcG, HKJ and JSL. RP and JRE performed colocalization analysis. AAP and OMS performed surface spike, ACE2 Fc binding assay and provided essential reagents. JRE, KO, JLH and AEF supervised the research. JRE, HS and RP wrote the manuscript and made the figures.

## Data and materials availability

all data are available in the manuscript or the supplementary materials.

## Supplemental Figures

**Figure S1**

(A) Mock infected differentiated nasal primary human airway epithelial (HAE) cells were infected and embedded to OCT. Cryostat sections were stained for spike (green), tetherin (red), phalloidin (grey) and DAPI (blue).

(B) Differentiated nasal primary human airway epithelial (HAE) cells were infected with SARS-CoV-2 (MOI 0.01). Cells were fixed at 48 hpi and embedded to OCT. Cryostat sections were collected and prepared for confocal microscopy. Sections were immunolabelled with antibodies against SARS-CoV-2 spike (green) - to reveal SARS-CoV-2 infected cells, tetherin (red), and with phalloidin-647 (grey) and DAPI (blue).

(C) SARS-CoV-2 infected HeLa+ACE2 cells (MOI 0.5) were fixed at 24 hpi and stained for spike (green) and tetherin (red). Infected cells display reduced tetherin levels, and broad loss of tetherin from the plasma membrane. Where cell surface tetherin remains, it is often clustered with spike staining (enlarged, arrows). Uninfected cells shown with asterisk.

(D) HeLa+ACE2 cells were infected with SARS-CoV-2 (MOI 0.5) and fixed at 24 hpi. Cells were stained for spike (green) and beta2microglobulin (red). No differences in beta2microglobulin were observed between infected and uninfected cells.

**Figure S2**

(A) Tetherin expression is induced with IFNα in A549 cells. Mock and IFNα treated (1000 U/mL, 24 hours) A549 cells were fixed and stained for tetherin (red) and analysed by confocal immunofluorescence microscopy.

(B) Additional electron micrographs of SARS-CoV-2 virions in infected T84 cells. T84 cells were infected with SARS-CoV-2 (MOI 0.5) and fixed at 24 hpi. Tethered virions were frequently present at the plasma membrane, and in intracellular compartments.

**Figure S3**

**(A)** Representative confocal immunofluorescence images of HeLa cells transiently transfected with SARS-CoV-1 ORF7a-FLAG or SARS-CoV-2 ORF7a-FLAG. SARS-CoV-1 ORF7a-FLAG predominately colocalizes with TGN46 (red), while SARS-CoV-2 ORF7a-FLAG shows additional staining outside that colocalizing with TGN46 (arrowheads).

**(B)** Manders’ coefficients were calculated to measure the ORF7a-FLAG overlap with TGN46. At least 54 cells per condition from three independent experiments were analysed. All data points, mean and standard deviation are plotted. Two-tailed, unpaired t-tests were performed. **** p < 0.0001.

**(C)** SARS-CoV-1 ORF7a-FLAG and SARS-CoV-2 ORF7a-FLAG stable cell lines were labelled with antibodies against FLAG (green), or the trans-Golgi marker TGN46 (red)

**(D)** As in (C) but with the cis-Golgi marker ZFPL1 (red).

**(E)** SARS-CoV-2 infected HeLa+ACE2 cells display fragmentation of Golgi markers. HeLa+ACE2 cells were infected with SARS-CoV-2 (MOI 0.5) and fixed at 24 hpi. Infected cells were identified by Spike staining (green) and cells were costained with Golgi markers TGN46 (top) and ZFPL1 (below). Areas of TGN46 and ZFPL1 are enlarged (right) to highlight Golgi fragmentation in SARS-CoV-2 infected cells.

**Figure S4**

(A) HeLa cells were transiently transfected with plasmids encoding ss-HA-Spike. 48 hours post transfection, cells were fixed and stained with antibodies against anti-HA (green) and anti-tetherin (red), and DAPI (blue).

(B) HEK-293T cells were transfected with the indicated constructs and their surface levels determined by flow cytometry using an antibody directed against the S2 subunit of the Spike protein. The light grey trace shows the non-transfected, unstained control and the dark grey trace the sample expressing the Spike construct. The mean fluorescent intensity was calculated for each sample and plotted. Error bars show the standard deviation for three biological repeats.

(C) HEK-293T cells were transfected with the indicated constructs and incubated with tissue culture supernatant containing soluble ACE2 (ACE2-Fc) and the amount of ACE2 binding determined by flow cytometry. The light grey trace shows the non-transfected control sample and the dark grey trace the sample expressing the Spike construct. The mean fluorescent intensity was calculated for each sample and plotted. Error bars show the standard deviation for three biological repeats.

(D) TetOne ss-HA-Spike cells were incubated with Doxycycline for 48 hours. Cells were fixed and stained for HA (green), tetherin (red) and with DAPI (blue) to show the presence of multinucleated cells.

(E) Flow cytometry was performed on mock or Doxycycline treated (48 hour) TetOne ss-HA-Spike cells to analyse surface CD71 levels.’

**Figure S5**

(A) SARS-CoV-2 ORFs were transiently expressed in HeLa cells. Cells were fixed, permeabilized and stained with anti-Strep antibodies and analysed by flow cytometry to confirm cells expressing strep-tagged SARS-CoV-2 ORFs.

(B) Representative confocal immunofluorescence microscopy images of fixed HeLa cells transiently transfected with SARS-CoV-2 ORFs. Anti-Strep (green), anti-tetherin (red), DAPI (blue).

**Figure S6**

(A) Intracellular flow cytometry was performed on mock HeLa and transiently transfected ORF3a-Strep HeLa cells to identify Strep positive (transfected) cells. The intracellular levels of tetherin were compared between the Strep negative (untransfected) and Strep positive (transfected) cells.

(B) Confocal immunofluorescence microscopy was performed on mock or ORF3a- Strep transfected cells to analyse the distribution of markers CIMPR, VPS35 and CD63.

(C) Confocal immunofluorescence microscopy was performed on mock or ORF3a- Strep transiently transfected cells to analyse the distribution of the early and late endolysosomal markers, EEA1 and Cathepsin D. Colocalisation between EEA1 and Cathepsin D could only be observed in ORF3a-Strep transfected cells. Enlarged area shows Cathepsin D within EEA1-positive compartments.

(D) Transmission electron microscopy was performed on mock or ORF3a-Strep transiently transfected HeLa cells. Endosomes and lysosomes appeared enlarged and containing non-resolved content in ORF3a-Strep transfected cells.

(E) Antibody uptake experiments were performed to follow the fate of endocytosed tetherin. ORF3a-Strep transient transfections were performed 48 hours prior to uptake experiments. Anti-tetherin antibodies were bound to live cells on ice for 30 minutes before a 2 hour chase at 37°C. Cells were fixed and immunolabelled using anti-LAMP1 (green), secondary anti-rabbit555 (red) antibodies, and DAPI (blue). Representative images are shown.

(F) Antibody uptake experiments were performed using anti-CIMPR antibodies. ORF3a-Strep transient transfections were performed 48 hours prior to uptake experiments. Anti-CIMPR antibodies (2G11 clone) were bound to live cells on ice for 30 minutes before a 2 hour chase at 37°C. Cells were fixed and

