## Supplementary figures and images for "Tetherin antagonism by SARS-CoV-2 enhances virus release: multiple mechanisms including ORF3a-mediated defective retrograde traffic"

### Supplemental Figure 1

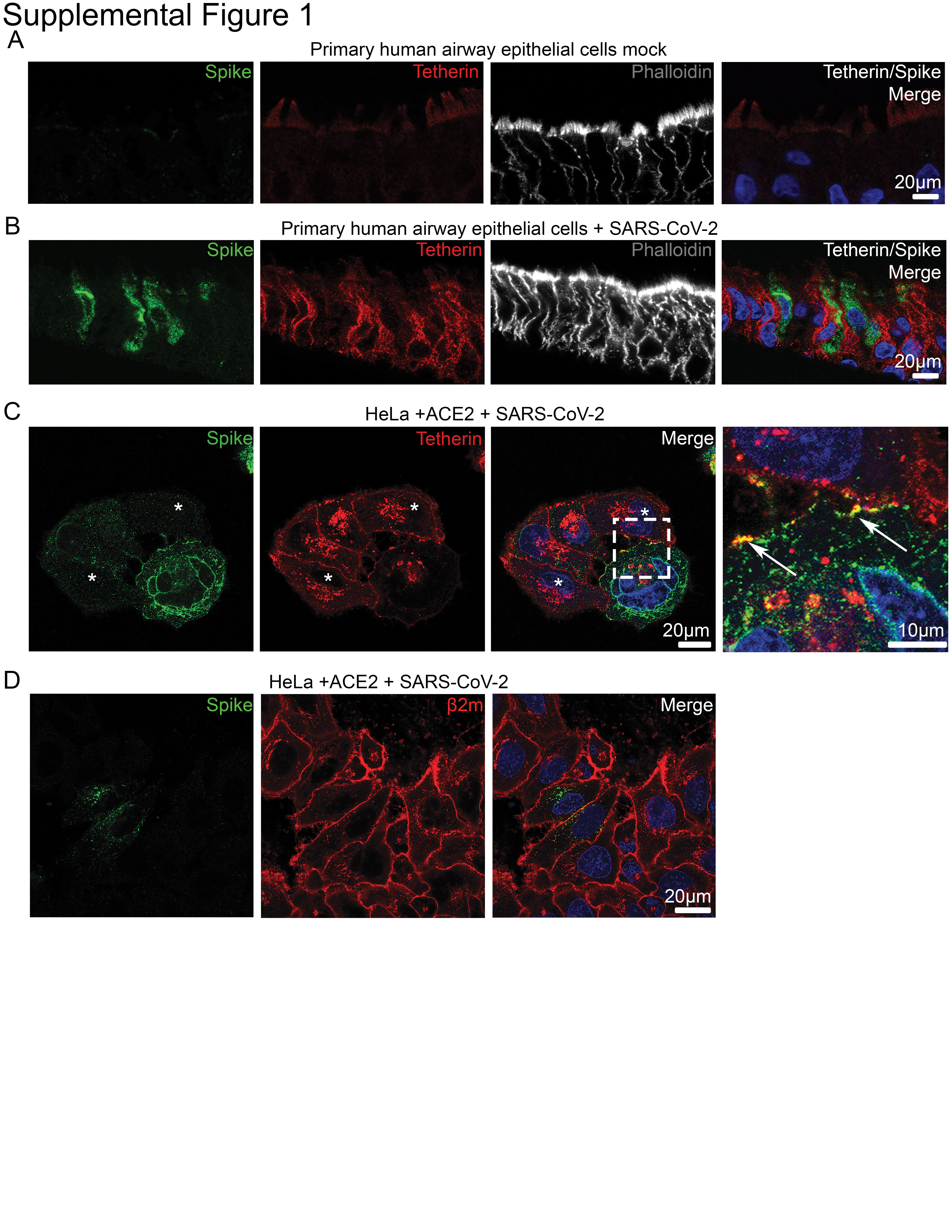

### Supplemental Figure 2

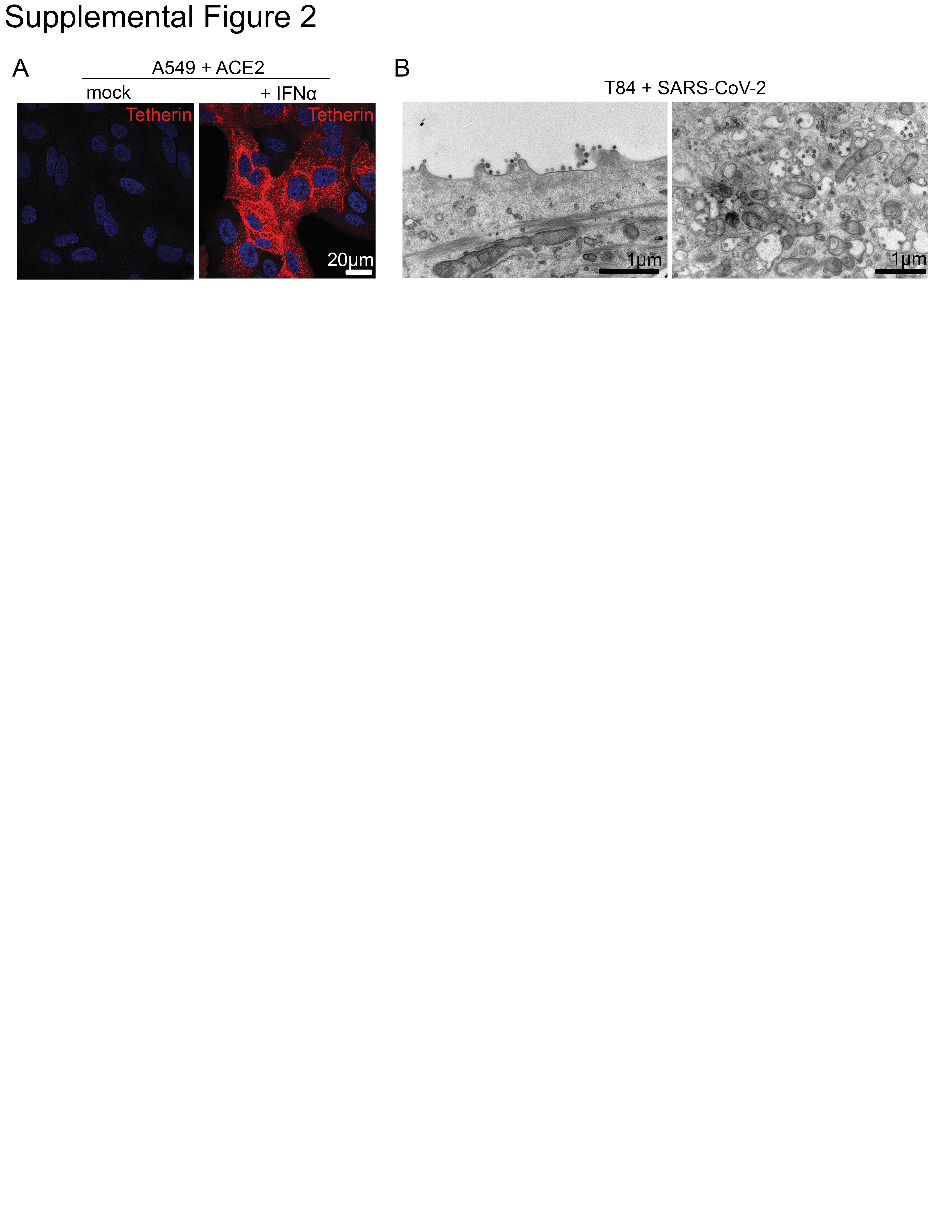

### Supplemental Figure 3

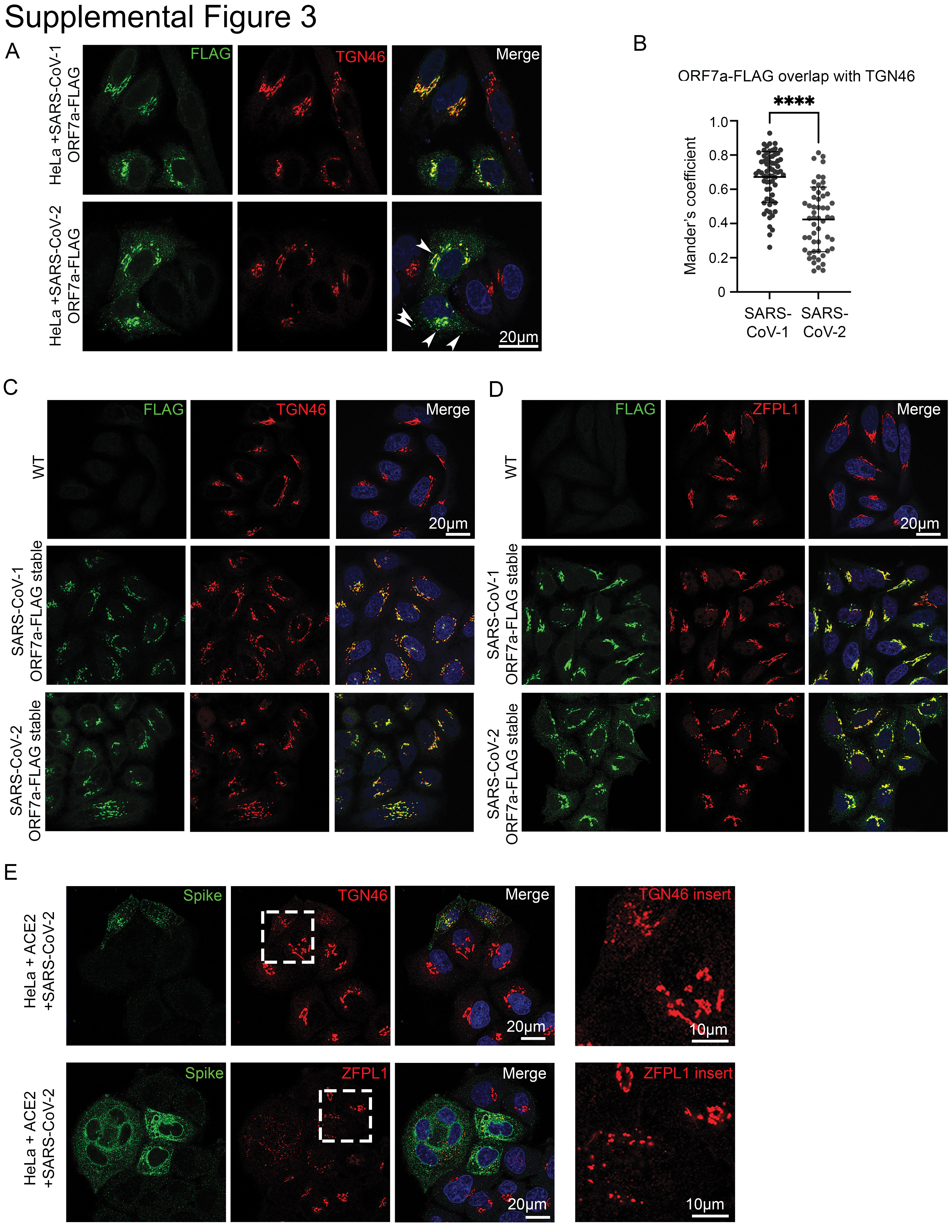

### Supplemental Figure 4

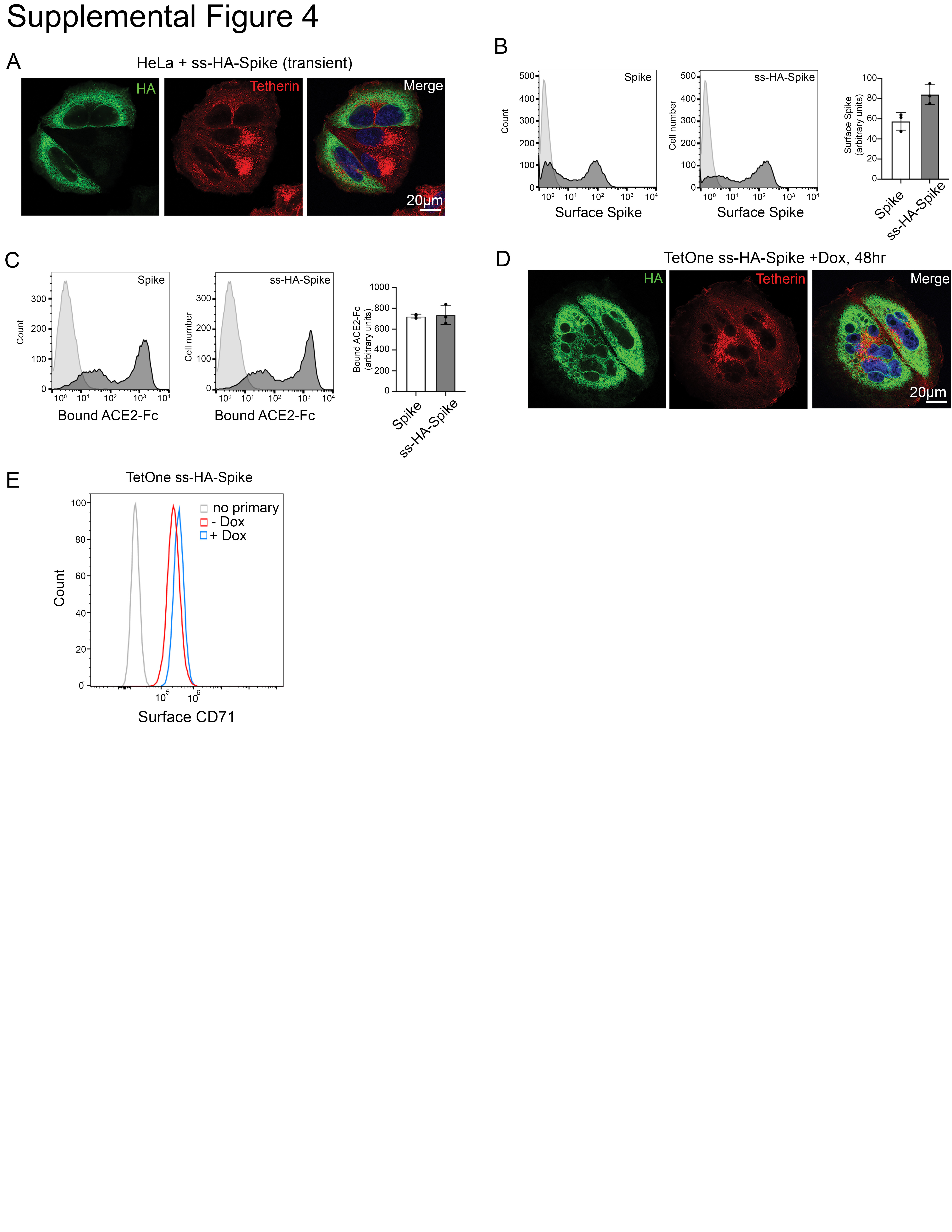

### Supplemental Figure 5

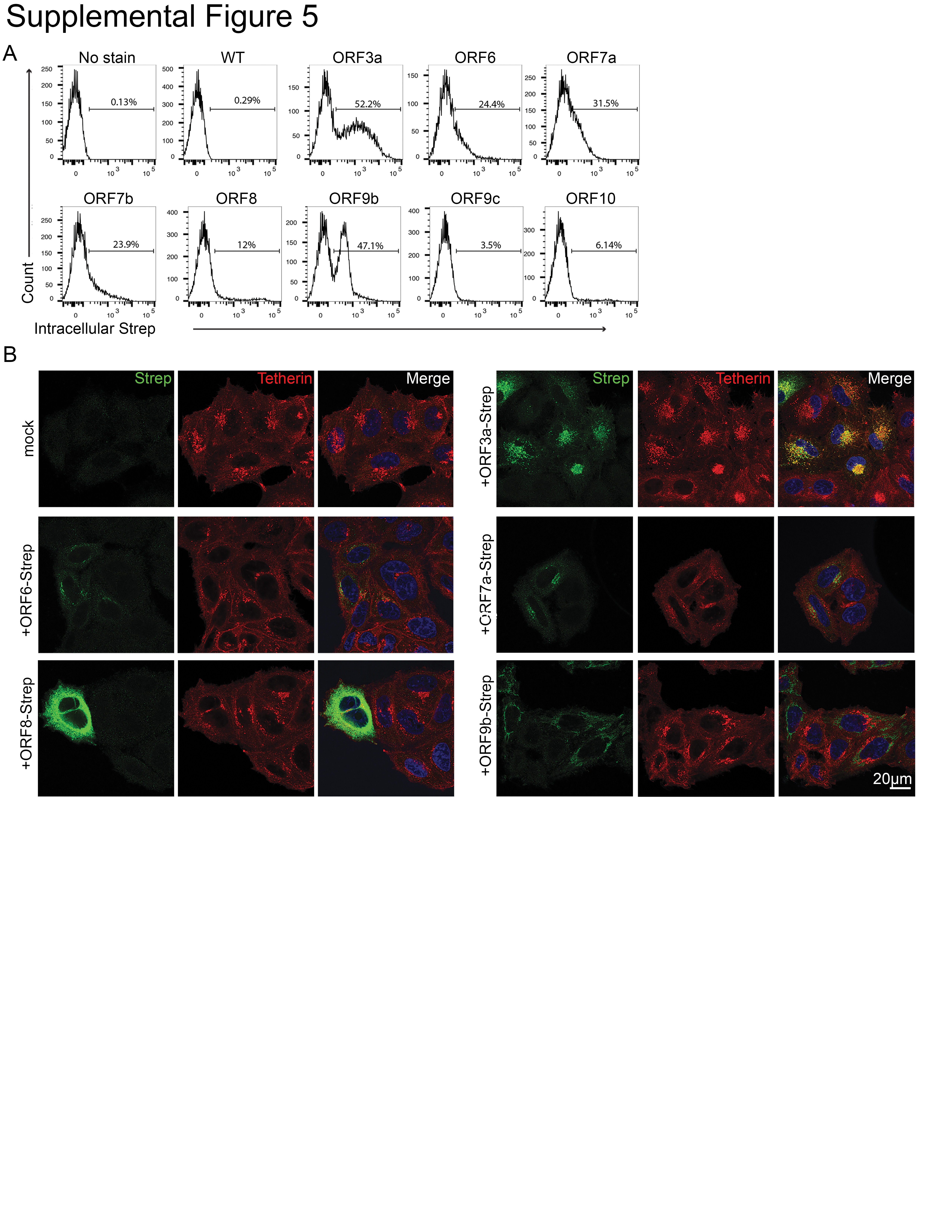

### Supplemental Figure 6

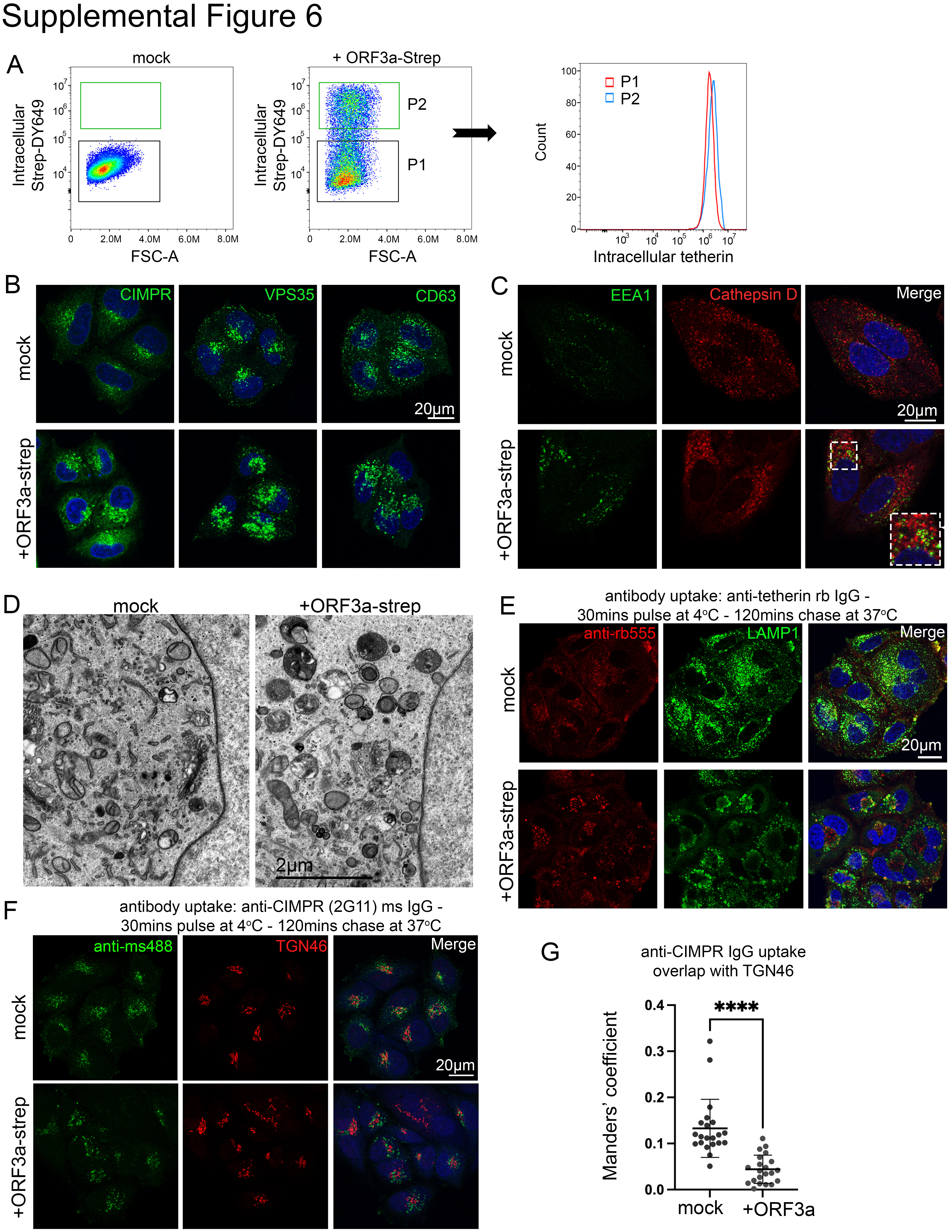
